# Development of a Cloud-Based IoT System for Livestock Health Monitoring Using AWS and Python

**DOI:** 10.1101/2024.06.08.598087

**Authors:** Harini Shree Bhaskaran, Miriam Gordon, Suresh Neethirajan

## Abstract

The agriculture industry is currently facing significant challenges in effectively monitoring the health of livestock. Traditional methods of health monitoring are often labor-intensive, inefficient, and insufficiently responsive to the needs of modern farming. As the number of IoT devices in agriculture proliferates, issues of scalability and computational load have become prominent, necessitating efficient and scalable solutions. This research introduces a cloud-based architecture aimed at enhancing livestock health monitoring. This system is designed to track critical health indicators such as movement patterns, body temperature, and heart rate, utilizing AWS for robust data handling and Python for data processing and real-time analytics. The proposed system incorporates Narrow Band IoT (Nb IoT) technology, which is optimized for low-bandwidth, long-range communication, making it suitable for rural and remote farming locations. The architecture’s scalability allows for the effective management of varying numbers of IoT devices, which is essential for adapting to changing herd sizes and farm scales.

Preliminary experiments conducted to assess the system’s performance have demonstrated its durability and effectiveness, indicating a successful integration of AWS IoT Cloud services with the deployed IoT devices. Furthermore, the study explores the implementation of predictive analytics to facilitate proactive health management in livestock. By predicting potential health issues before they become apparent, the system can offer significant improvements in animal welfare and farm efficiency. The integration of cloud computing and IoT not only meets the growing technological needs of modern agriculture but also sets a new benchmark in the development of sustainable farming practices. The findings from this research could have broad implications for the future of livestock management, potentially leading to widespread adoption of technology-driven health monitoring systems in agriculture. This would help in optimizing the health management of livestock globally, thereby enhancing productivity and sustainability in the agricultural sector.

## 1. Introduction

The Canadian agricultural sector, particularly in Nova Scotia, is significantly driven by animal husbandry, which not only supports the livelihoods of farm families and sustains rural communities but also enriches cultural diversity and promotes social cohesion. This sector is a cornerstone of the provincial economy, evidenced by the steady growth in GDP contributions from agriculture, forestry, fishing, hunting, and related industries over the past decade. Notably, aquaculture and animal production have seen increases of 17.9% and 3.2%, respectively, underpinning their importance to the economic landscape of Nova Scotia [1].

The Livestock Health Monitoring (LHM) systems are crucial in this context, where sustainability, agility, and precision are paramount. As of 2023, the dairy sector alone in Nova Scotia generated $33.5 million, underscoring the necessity for advanced technological solutions to optimize productivity and profitability [2].

Historically, livestock health monitoring has relied on intermittent manual observations, which often result in delayed responses to health issues and inefficient farm management. The advent of integrated modern technologies, including IoT and cloud computing, offers transformative potential for real-time, proactive animal health monitoring. This shift not only promises to enhance animal welfare but also to improve the operational efficiency of farms [3, 4].

This research is positioned to further these advancements by proposing a scalable Cloud-based IoT architecture aimed at refining the precision and efficiency of real-time livestock health monitoring. Incorporating technologies such as Narrowband IoT (Nb IoT) enhances connectivity, extends battery life, and ensures robust data transmission even in remote settings, which is crucial for extensive farming operations [5].

### 1.1. Technological Innovations and Challenges of Livestock Farming

Traditional methods for livestock health monitoring are labor-intensive and often inefficient. For example, behavioral observations critical for managing reproductive health can consume up to 30% of labor resources on commercial farms [6, 7]. These methods underscore significant inefficiencies in resource allocation and highlight the critical need for automated and precise monitoring systems that can significantly reduce labor costs and improve health detection rates [8].

Precision Livestock Farming (PLF) tools represent a leap forward in managing animal health. These systems utilize advanced sensors and data analytics to provide real-time insights into animal behavior and health. PLF tools help overcome the limitations of human observation, facilitating early detection of health issues and more efficient herd management [9].

As the global population is projected to reach 9.7 billion by 2050, the demand for animal products is expected to surge, necessitating a 70% increase in global food production, with meat demand alone expected to rise by over 50% [10, 11]. The livestock industry’s expansion has been rapid, fueled by globalization and increasing consumer demands across diverse markets. However, this growth brings challenges concerning sustainability, environmental impacts, and animal welfare, making it imperative to balance increasing meat production with responsible farming practices to ensure long-term industry viability [12–15].

### 1.2. IoT and Sensor Technologies in Livestock Health Management

The IoT framework for LHM integrates various components essential for effective health monitoring. Sensors like accelerometers and temperature sensors are deployed extensively to monitor vital physiological parameters such as heart rates and body temperatures, which are indicative of the animals’ general health. These sensors provide a continuous stream of data, which is crucial for timely interventions. Moreover, communication technologies such as ZigBee, RFID, Bluetooth, and Wi-Fi play pivotal roles in the efficient transmission of this data [16].

Despite the advancements, certain limitations persist with technologies like ZigBee, which currently supports monitoring only one animal at a time. In contrast, integrating LoRaWAN and NbIoT has shown promise in enhancing data transmission efficiency, demonstrating low operational costs and extended communication ranges, even in complex environments [41]. The amalgamation of NB-IoT and LoRa technologies is particularly advantageous, offering extended transmission distances and reduced operational costs. These features are essential for large-scale farm applications, where traditional cellular communications may prove less energy-efficient and more costly [17, 18].

Additionally, incorporating cloud services in livestock health monitoring provides enhanced data management, real-time analysis, scalability, and improved decision-making. Continuous collection and analysis of health data from IoT sensor can detect health issues early, enabling timely interventions. Immediate alerts about abnormal conditions such as changes in body temperature, heart rate, or activity levels are sent to farmers and veterinarians to respond quickly to potential health problems.

Moreover, cloud platforms provide scalable framework that can handle huge volumes of data from numerous devices. Hence data from a large farm can swiftly scale their data storage and processing capabilities up or down depending on the requirement without significant investment in physical hardware.

### 1.3. Gaps in the Current Knowledge Landscape

The adoption of advanced technologies in livestock health monitoring on small-scale farms faces specific challenges. Research focusing on these barriers and developing tailored solutions is crucial for wider technology adoption [19]. While the immediate benefits of technology adoption in LHM are evident, there is a notable gap in literature concerning the long-term impacts on livestock health, farm sustainability, and economic viability. Longitudinal studies are required to provide deeper insights into these aspects [20]. The ethical implications of deploying advanced technologies in livestock monitoring remain under-explored. Issues such as data privacy, animal welfare, and societal impacts need comprehensive examination to ensure responsible technology implementation [21]. The existing literature often focuses on specific regions or countries, lacking a global perspective that accounts for varied agricultural practices, socio-economic factors, and environmental conditions. Developing universally applicable livestock health monitoring strategies requires a broader understanding of these global dynamics [22]. **Supplementary Table S1** captures various IoT applications, objectives, sensors, and technologies used in livestock monitoring from recent studies.

The overall objectives of the study are,

1. To develop a Cloud-based IoT architecture that enhances real-time monitoring and management of livestock health.
2. To create a collaborative web-based platform that serves as a nexus for interaction among farmers, researchers, and regulatory bodies, fostering a community of practice that enhances knowledge exchange and supports sector-wide improvements.
3. To utilize advanced data analytics, powered by Python, for robust data processing and interpretation, enabling precise health assessments and predictive analytics for proactive farm management.

### 1.4. Cloud-Based System in Animal Husbandry

The goal of smart animal farming is to harness the potential of cloud computing technology and internet to give a boom in the pasturage [52]. Cloud technology offers a network of remote servers hosted on the internet for the purpose of storing, managing, and processing massive volumes of data to facilitate data-driven farming. This ability to store and manage huge volumes of data, cost effectiveness and remote accessibility makes it optimal solution for addressing the challenges faced by the agricultural sector [10]. A Livestock monitoring system is a cutting-edge solution architected and developed using sensors, GPS, etc., and integrating all these with a network protocol for communication. This monitoring system enables farmers to remotely check their farms and take actions immediately. Amazon AWS for Cloud Deployment in Livestock Health Monitoring Leveraging Amazon AWS for cloud deployment in livestock health monitoring provides several critical advantages. AWS offers unparalleled scalability, allowing the system to handle data from numerous IoT devices without performance degradation, even as the number of devices increases [53]. This flexibility is crucial for adapting to varying farm sizes and the dynamic nature of livestock operations. AWS ensures high availability and reliability, with services distributed across multiple geographic regions and availability zones. This redundancy minimizes the risk of downtime and data loss [54], ensuring continuous monitoring and prompt responses to any health issues in the livestock.

Security is a paramount concern in any cloud deployment, and AWS provides robust security features, including encryption, access control, and continuous monitoring to safeguard sensitive data [55]. Compliance with industry standards and regulations is also easier with AWS’s comprehensive suite of compliance certifications. Cost efficiency is another significant benefit. AWS’s pay-as-you-go pricing model [56] allows for cost-effective scaling, reducing the need for large upfront investments in physical infrastructure. This model ensures that resources are used efficiently, and costs align with actual usage, which is particularly beneficial for small and medium-sized farms.

Integration capabilities are a strong suit of AWS, facilitating seamless interaction between various services such as AWS IoT Core, DynamoDB, and Amazon Pinpoint. This integration enables the creation of a cohesive and efficient system that supports real-time data processing, storage, and communication. Moreover, AWS’s extensive global infrastructure provides low-latency access and fast data processing, crucial for real-time health monitoring applications where timely interventions can significantly impact animal welfare.

### 1.5. Livestock Health Monitoring

LHM is of utmost importance when overall farm management is considered. It is essential to maintain proper check of the livestock since good health and well-being of livestock are mandatory for sustainable production of milk [57]. It also paves way for farmers to have a check on the animal behaviour, identifying early signs of disease detection [58]. Timely intervention helps prevent the spread of diseases within the herd. This not only saves individual animals but also the whole livestock population. In addition to this, selecting animals with desirable health traits for breeding can help ensure that the best qualities are passed on to future generations [59]. Another important reason is that quality and safety of food to consumers [60]. Healthy livestock produce high-quality products which is highly essential for maintaining consumer demands. It avoids diseases spreading from animals to humans [61].

Furthermore, by monitoring key health metrics farmers can optimize breeding programs, selecting healthier animals with desirable traits to improve the overall herd’s genetics [62]. Besides, this health monitoring also helps understanding the movement pattern such as grazing, rumination and idling behaviours [63].

### 1.6. Narrow-Band Internet of Things

Nb-IoT is a wireless IoT that works on the principle of Low Power Wide Area Networks (LPWANs) technology [64], providing a greater volume of devices to share bandwidth than traditional cellular networks like 2G, 3G, and 4G. The unused frequencies within a carrier’s licensed bands are utilized by these devices, thereby consuming less power than many other types of networks. Since the NB-IoT networks use narrower frequency band, it allows a larger volume of devices to occupy one of the network’s “cells.” This enhances coverage by repeating transmissions and increasing the receiver’s ability to resolve messages. Advantages of using Nb-IoT includes Low power consumption; Power Saving Mode (PSM); Extended Discontinuous Reception (eDRX); Efficient spectrum used; Ubiquitous Coverage and Connectivity; Low cost; Strong signals; Good battery life [65].

### 1.7. Agile Methodology

The proposed research follows the principles of Agile Methodology that has dynamic phases called sprints. In contrast to other models in software development, Agile method is flexible, efficient and follows an iterative approach thus ensuring requirement satisfaction [66]. The various phases of Agile method are elaborated in the subsequent paragraphs.

#### 1.7.1 Requirements

The primary step of any successful architecture lies in the ideation stage. This serves as a foundation to build the LHM system pertaining to the satisfy the following parameters. The LHM model that is based on AWS Cloud architecture and IoT must be a very secure system as it stores and monitors private data. Hence this should prevent breaching data through unauthorised access. Any robust model must be handled easily and efficiently without much human interference. This architecture complies to this as troubleshooting takes place to isolate fault thereby not breaking the operation of the entire system. It should meet the changing demand by allocating or removing resources depending on the incoming traffic from the system. The model is flexible to changing herd size thereby not incurring more cost. The architecture must be able to run through its complete lifecycle. All the desired services and components must be available 24/7.

#### 1.7.2 Planning

A crucial step in modelling this architecture, is the planning phase which involves deploying the technological stack for the application. The various features to be included in the LHM is outlined as follows:

Feature 1: A secure model that prevents unauthorised access.

Feature 2: Model that works seamlessly irrespective of resources being coupled or decoupled.

Feature 3: The system must be capable of handling increases or decreases in data load efficiently.

Feature 4: Must allow more than 500 IoT sensors to operate simultaneously.

Feature 5: The framework must be accessible in remote areas.

Feature 6: Identify any anomalies in the behaviour of animals.

Feature 7: Perform analytics and create visualizations from the collected data.

Feature 8: SMS notification to farmers and concerned authorities, vet.

Feature 9: Record and store the event. Feature 10: Distributed storage.

#### 1.7.3 Designing

Any cloud-based model must comply to the five pillars of well-architected framework to have satisfied the requirements of the proposed work. This involves the framework to perform the necessary actions as coded and must be flexible to changes. The system must anticipate failures, learn, and recover from it. The proposed model must have a strong secure system to protect the data when in transit or at rest. It should also anticipate any security events Ensure that the proposed work functions effectively and efficiently as expected. Correct usage of resources. It must be cost-efficient and avoid unnecessary expenditure.

#### 1.7.4 Developing

The technology stack is determined based on the features that are necessary to build the LHM. This includes a variety of AWS Services, software tools and technologies that are deployed during the development phase. The overall performance of a model depends on the efficiency of the components selected and their effective utilization. Hence choosing the right technology infrastructure is of utmost importance to achieve desired results. Various AWS Services, software tools, and technology used in developing this model are outlined as follows: To satisfy the requirements mentioned in step 1, the proposed model is built on AWS Cloud infrastructure as it provides secured access management, and cost-effective and flexible resource allocation [67]. In addition, it also offers a well-developed edge computed service that allows resources to be coupled or decoupled alongside the other required IoT services. Python is a high-level programming language that offers high readability, supports multiple programming paradigm with a vast number of libraries [68]. It is essential in this framework as it has an excellent SDK for developing cloud-based services [69]. This is utilized for the development of software deployed in IoT devices, and AWS Lambda functions, which are used as data processing triggers for AWS data pipelines. The smart livestock health monitoring architecture built in AWS combines the various AWS serverless services that perform individual and independent tasks.

#### 1.7.5 Testing

Data load testing ensures the system’s readiness to handle large data volumes continuously while maintaining acceptable levels of average latency and error counts. Data integrity testing verifies that data is correctly populated across all locations and that insertion events occur in the proper sequence. Functional testing guarantees proper data acceptance, processing, and retrieval under high loads. Stress testing evaluates the system’s durability, scalability, and performance. Security and access control testing ensures that only authorized users can access the ingested data.

In terms of evaluating results, test completion and success criteria are assessed by analyzing test logs and various performance metric charts.

#### 1.7.6 Deploying

During deployment, the architecture utilizes Terraform for provisioning resources in the AWS infrastructure. Hashicorp Terraform is employed to create, modify, and provision these resources using JSON configuration files. Commands are then executed to adjust the infrastructure according to requirements, with Terraform’s robust capabilities available to restore infrastructure state if necessary.

## 2. Materials and Methods

### 2.1. Ethical Declaration

The empirical data supporting this study was generously provided by my distinguished colleagues at Wageningen University & Research and is linked to an independent, prior experiment. The study in question was authorized by the Central Committee on Animal Experiments (CCD) and the Animal Experiments Department (IVD) of the Netherlands, ensuring that the ethical standards and optimal procedures were followed. The Department of Animal Sciences of the Care of Animals Used for Scientific Purposes (CARUS) at Wageningen University & Research carefully reviewed and approved any additional, non-invasive treatment of animals for the current study. For this investigation, the appropriate approval number is 20210521ADP [70].

#### 2.1.2 Study Design and Animal Housing: A Methodological Approach

50 male piglets (Tempo × Topigs Norsvin TN70), weighing an average of 25 kg and born about nine weeks prior to use, participated in the long-term study. Rooms 14 and 15 of Wageningen University & Research’s CARUS facility served as the home for these piglets. Six piglets, ranging in age from 86 to 108 days, were selected from this cohort to undergo a thorough examination of their behavioural and physiological adaptations. Acclimatization took place throughout the first week, giving the piglets time to become used to their new surroundings, food, and care routine. Measuring 2.86 by 1.16 meters, each pen was large enough to accommodate two piglets and was furnished with all the necessities, including toys. Lights were kept on from 7:00 to 19:00, and the room temperature was adjusted to suit the demands of the piglets. The piglets were fed a high-protein diet supplemented with dried fermentation solubles, designed to meet their specific growth rate requirements. Their diet also consisted of grain and hay. Feed was provided twice daily, once at 08:00 and once at 17:00, with water available ad libitum[71].

#### 2.1.3 Data Collection

Data used in this study is collected from already mounted Zephyr BioHarness belt application on a Topigs Norsvin TN70 piglet (**Figure 2a**) in a swine farm. The design structure, layout, and equipment of the sensor installed on the pig can be found in (**Figure 2b and 2c**) illustrate two components of a single system. This system automates the process of data collection and transmits these data to the system. The used approach represents a non-invasive method for data collection.

**Figure 2a).**
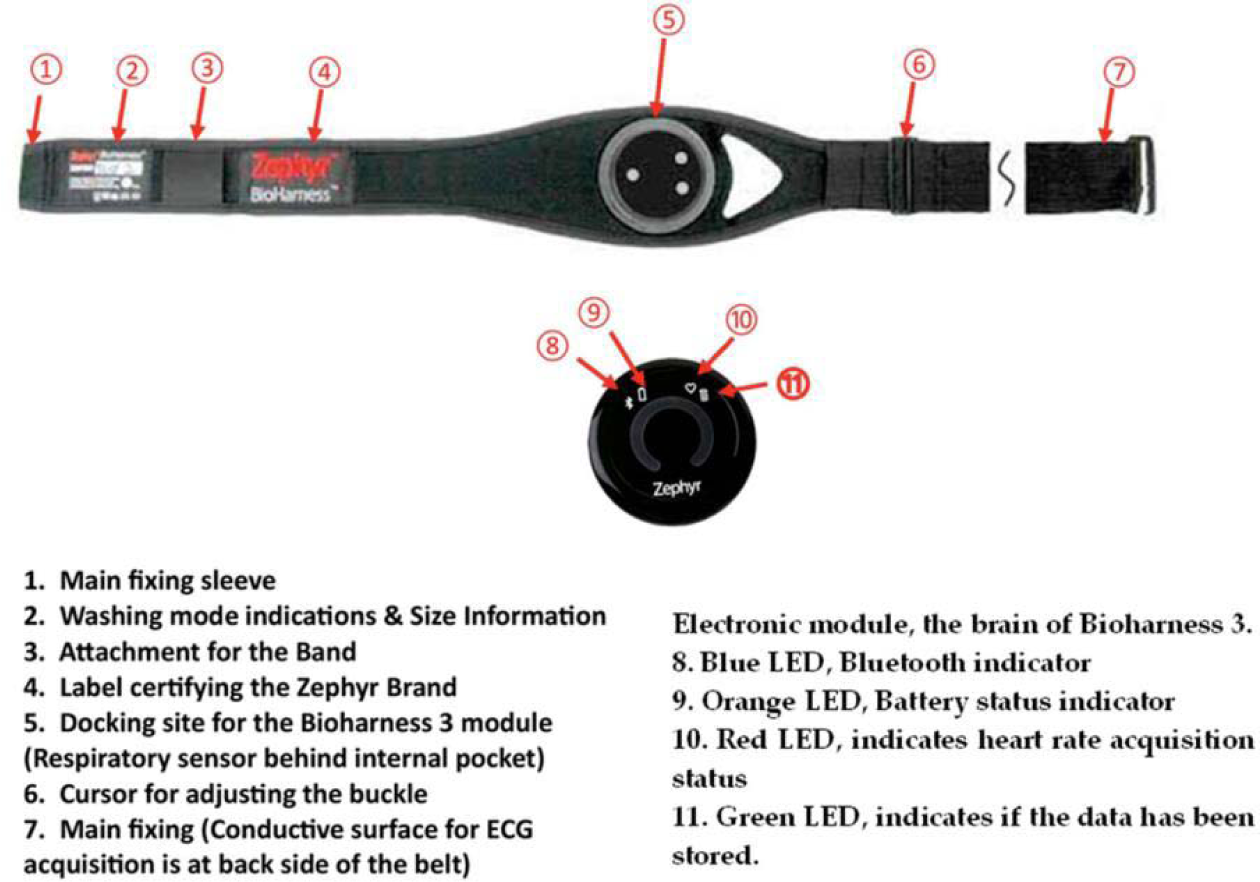
Photograph of the Zephyr BioHarness multimodal sensor platform used for data collection from Topig s Norsvin TN70 piglets.

**Figure 2b).**
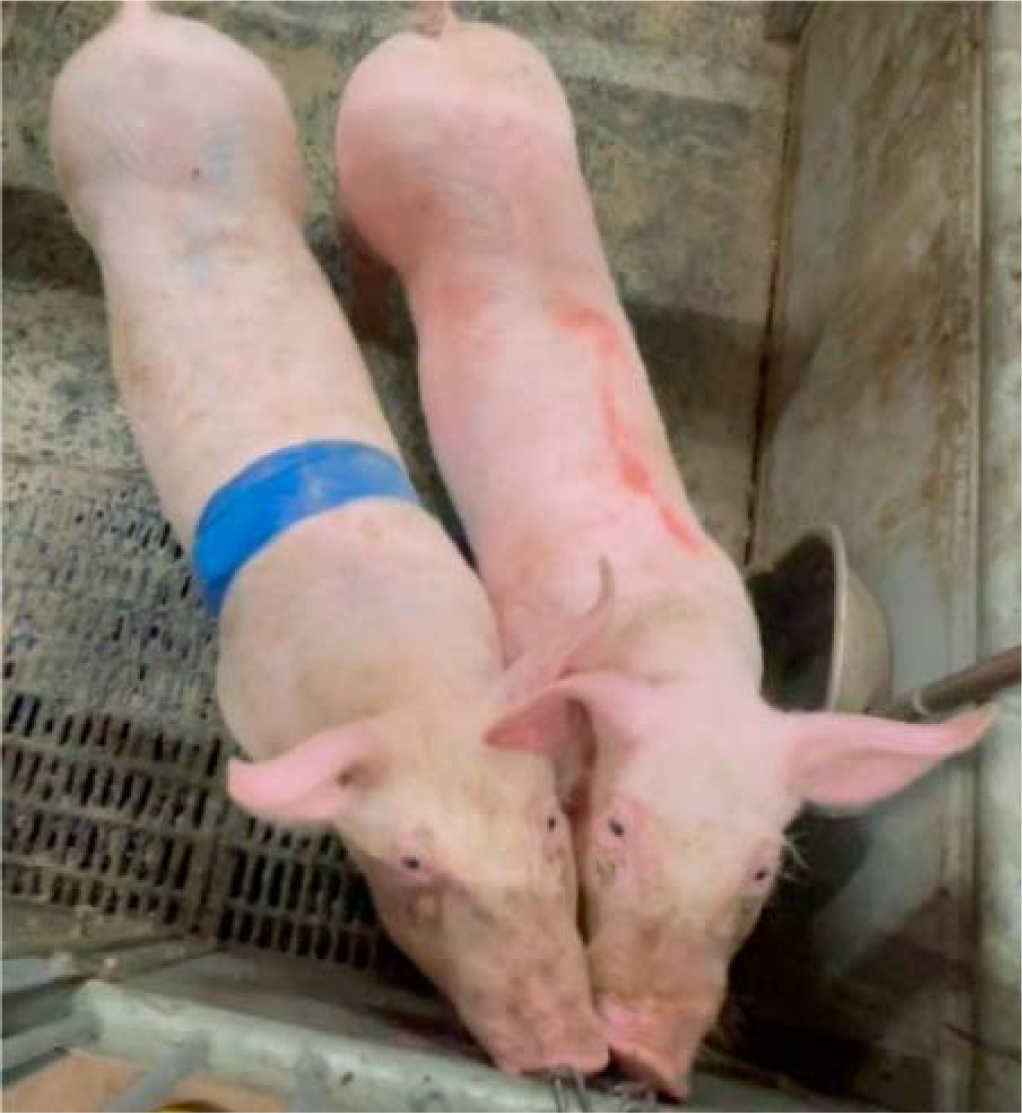
Overhead view portraying the Zephyr BioHarness 3.0 strap’s arrangement around the piglet’s chest, reinforced with a Vetrap bandage.

**Figure 2c).**
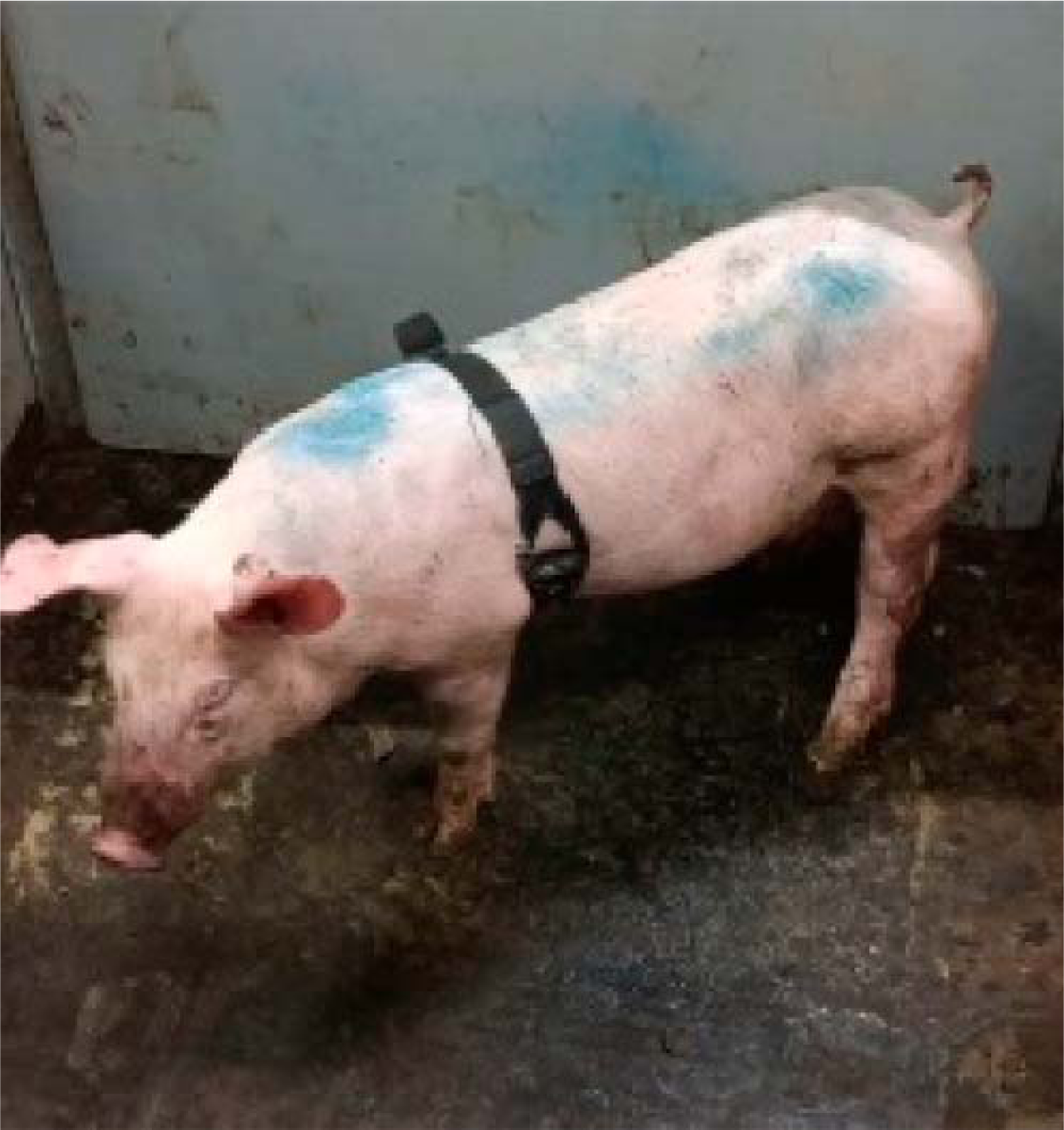
Side perspective showcasing the piglet wearing the Zephyr BioHarness.

#### 2.1.4 Workflow

The sensor data collected are in a raw format, hence it is necessary to analyze and format the data into usable form for developing a custom ML model. The methods, process, and approaches to understand how the data from IoT devices is processed and modeled and integrate this trained ML model into the production environment along with other AWS services [72].

The methodology being considered is to gather the unique characteristics of each pig, considering important health indicators. The collected dataset is primarily understood using domain knowledge to identify any issues or inconsistencies. Identifying and handling missing values in the dataset serves as the second step which is done using AWS S3. Depending on the nature of the missing data, the rows or columns containing missing values, NaN values are either removed or imputed using statistical methods (e.g., mean, median, mode). The dataset is then handled for any outliers and is uploaded to Sage Maker for making predictions. Data transformation techniques including one hot encoding are performed on relevant columns to transform categorical values to a usable form.

In the subsequent steps, unused or unnecessary columns are dropped, and predictions are performed. The dataset is then batch tested for forecasting the heart rate values in the future. This helps to accurately classify health status based on the animal’s specifics and its environment. It leads to improved early detection of potential health problems and prevention of disease outbreaks in the livestock industry.

The data acquired from the tripartite sensor is listed in **Table 1**. The collected data includes temperature, heart rate, breathing rate etc.

**Table 1.**
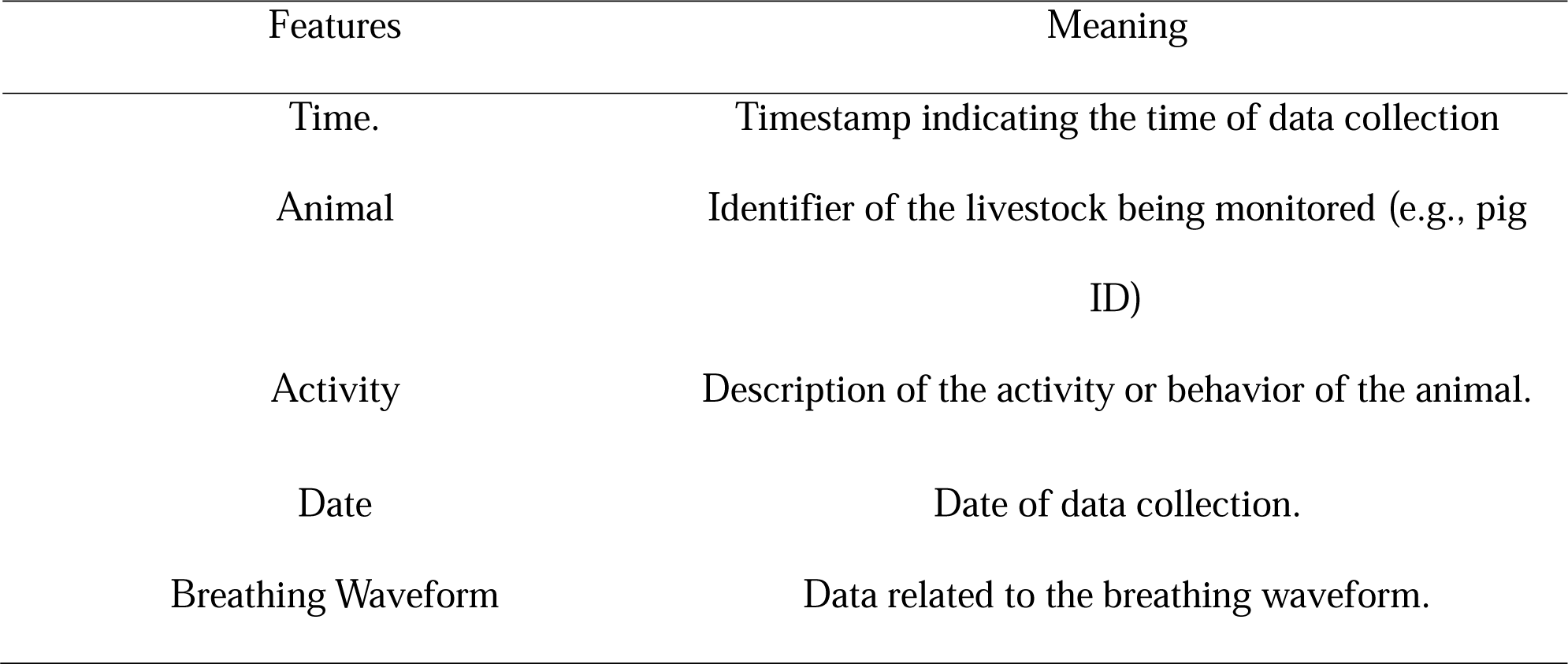
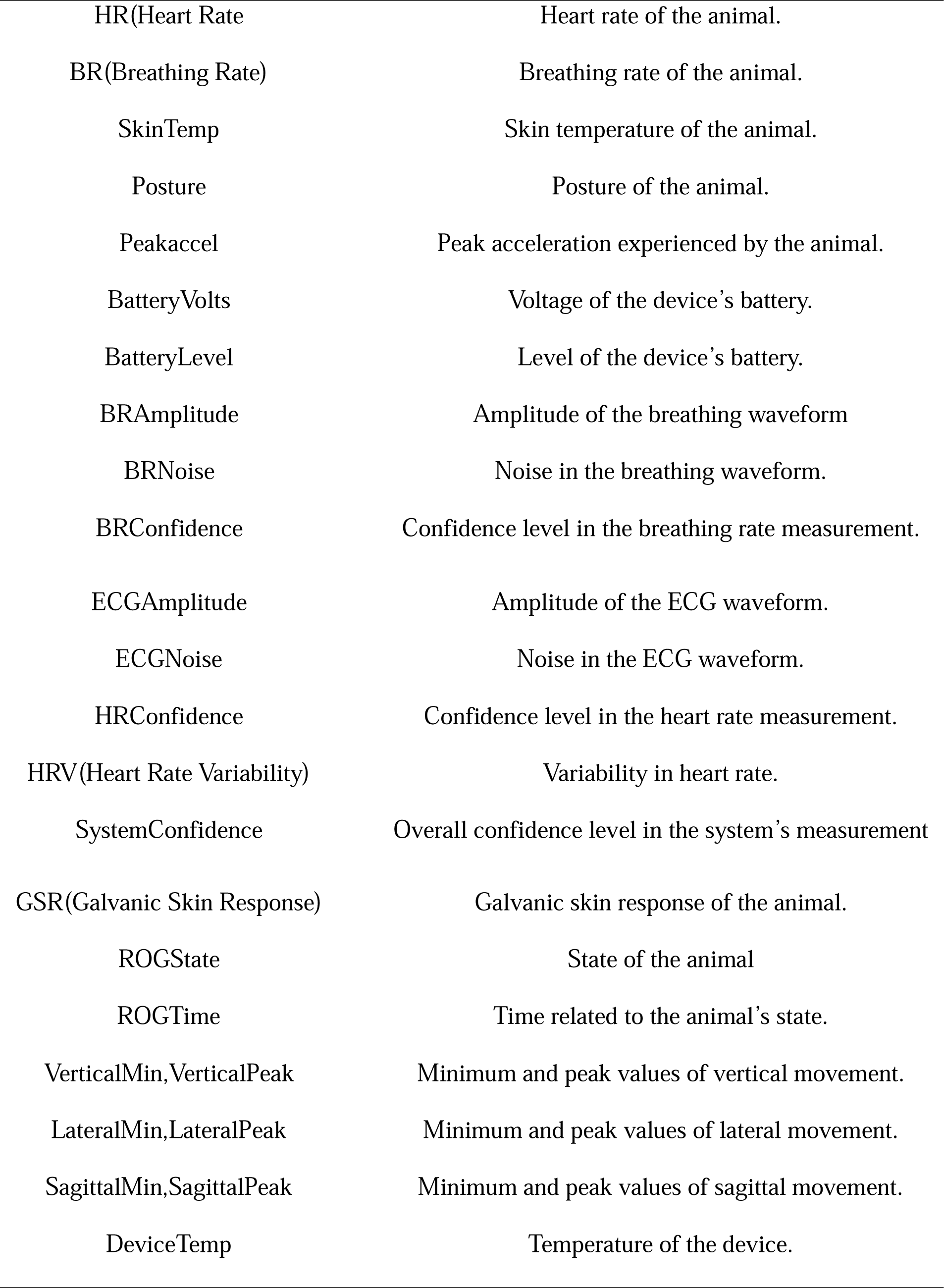
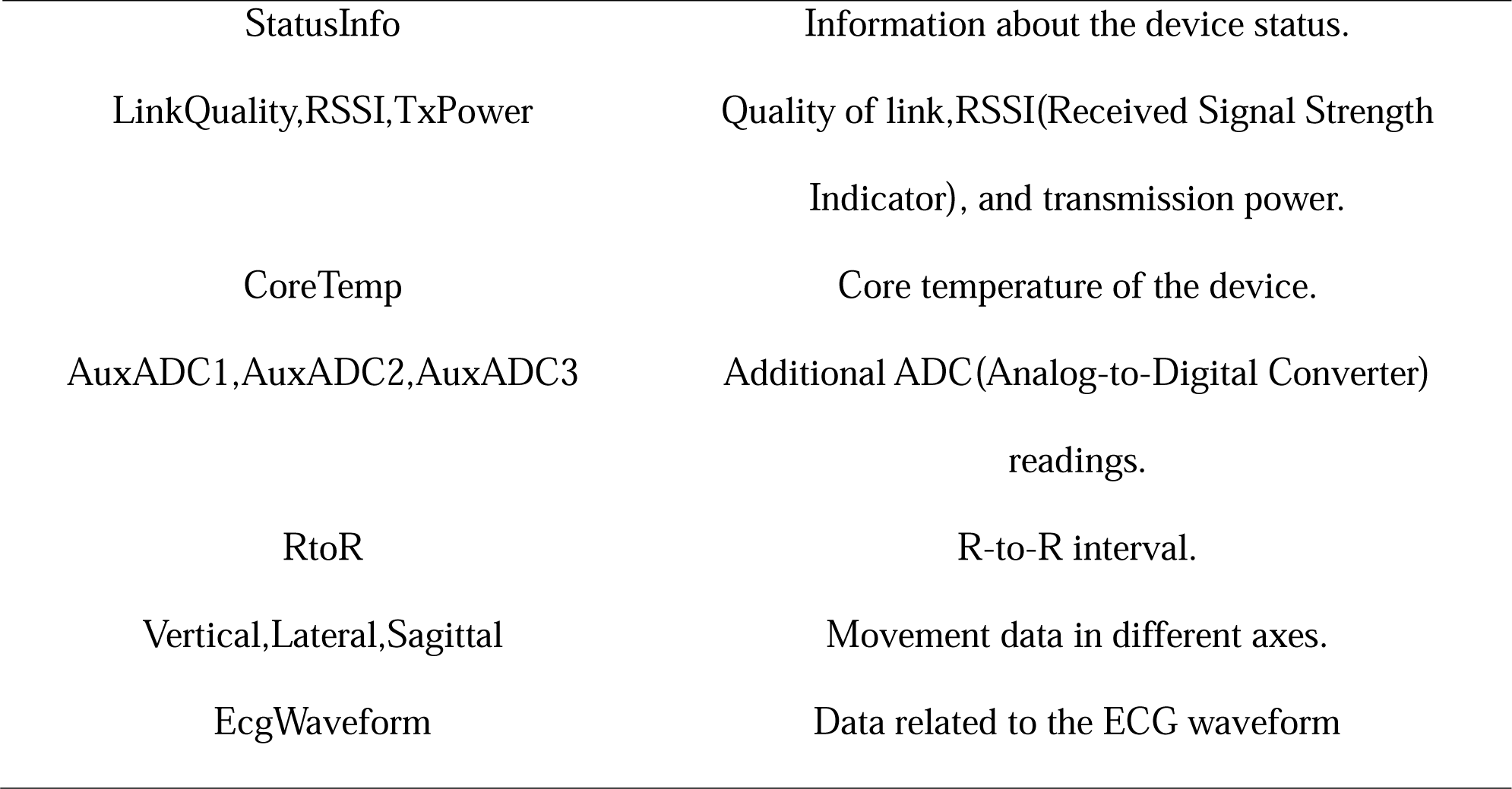
Various parameters measured using the tripartite sensor installed on the body of the pig.

| Features | Meaning |
| --- | --- |
| Time. | Timestamp indicating the time of data collection |
| Animal | Identifier of the livestock being monitored (e.g., pig ID) |
| Activity | Description of the activity or behavior of the animal. |
| Date | Date of data collection. |
| Breathing Waveform | Data related to the breathing waveform. |
| HR(Heart Rate | Heart rate of the animal. |
| BR(Breathing Rate) | Breathing rate of the animal. |
| SkinTemp | Skin temperature of the animal. |
| Posture | Posture of the animal. |
| Peakaccel | Peak acceleration experienced by the animal. |
| BatteryVolts | Voltage of the device's battery. |
| BatteryLevel | Level of the device's battery. |
| BRAmplitude | Amplitude of the breathing waveform |
| BRNoise | Noise in the breathing waveform. |
| BRConfidence | Confidence level in the breathing rate measurement. |
| ECGAmplitude | Amplitude of the ECG waveform. |
| ECGNoise | Noise in the ECG waveform. |
| HRConfidence | Confidence level in the heart rate measurement. |
| HRV(Heart Rate Variability) | Variability in heart rate. |
| SystemConfidence | Overall confidence level in the system's measurement |
| GSR(Galvanic Skin Response) | Galvanic skin response of the animal. |
| ROGState | State of the animal |
| ROGTime | Time related to the animal's state. |
| VerticalMin,VerticalPeak | Minimum and peak values of vertical movement. |
| LateralMin,LateralPeak | Minimum and peak values of lateral movement. |
| SagittalMin,SagittalPeak | Minimum and peak values of sagittal movement. |
| DeviceTemp | Temperature of the device. |
| StatusInfo | Information about the device status. |
| LinkQuality,RSSI,TxPower | Quality of link,RSSI(Received Signal Strength Indicator), and transmission power. |
| CoreTemp | Core temperature of the device. |
| AuxADC1,AuxADC2,AuxADC3 | Additional ADC(Analog-to-Digital Converter) readings. |
| RtoR | R-to-R interval. |
| Vertical,Lateral,Sagittal | Movement data in different axes. |
| EcgWaveform | Data related to the ECG waveform |

**Table 2.** List of AWS services required to satisfy the feature requirements outlined in Planning phase.

| Features | AWS Service |
| --- | --- |
| Feature 1 |  |
| Secure model that prevents unauthorised access | AWS Device Defender, |
| Feature 2 |  |
| Model that works seamlessly irrespective of resources being coupled or decoupled | AWS Greengrass |
| Feature 3 |  |
| It must handle increase or decrease in traffic | Auto Scaling |
| Feature 4 |  |
| Must allow more than 200 IoT sensors to | IoT Core Services |
| operate simultaneously |  |
| Feature 5 |  |
| The framework must be accessible in remote areas | Nb-IoT protocol |
| Feature 6 |  |
| Identify any anomalies in the behaviour of animals | AWS CloudWatch |
| Feature 7 |  |
| Perform analytics and create visualizations from the collected data | AWS SageMaker, Quicksight |
| Feature 8 |  |
| SMS notification to farmers and concerned authorities, vet | AWS SNS |
| Feature 9 |  |
| Record and store the event | DynamoDB |
| Feature 10 |  |
| Distributed Storage | AWS S3 |

## 3. Results

Technological assumptions for the proposed study include reliable network connectivity to transmit data from the farms. Adequate data storage for storing the data with proper encryption and access control to ensure data privacy. Other economic considerations include economically viable for the farm outweighing the costs. The solution is scalable, allowing for expansion as the livestock population grows or as additional farms are integrated into the system.

### 3.1 Services used in AWS Cloud

Using AWS serverless solutions, the proposed design for cattle farming in **Figure 3a** integrates a variety of services, encompassing development, computation, storage, databases, analytics, networking, mobility, management, IoT, business apps, and security. Essential components such as AWS IoT Core, Lambdas, DynamoDB, S3, Machine Learning, Notifications, Analytics, Logging, and User Identities were grouped to establish a scalable and resilient monitoring system. These groupings were organized to meet the functional requirements outlined in Phase 2 of the Materials and Methods section. Using requisite services enables the user to manage the current data and make future forecasts using historical data. Furthermore, Phase 3 of the Materials and Methods section delineates the architecture in alignment with the fundamental principles of a well-structured design.

**Figure 3a).**
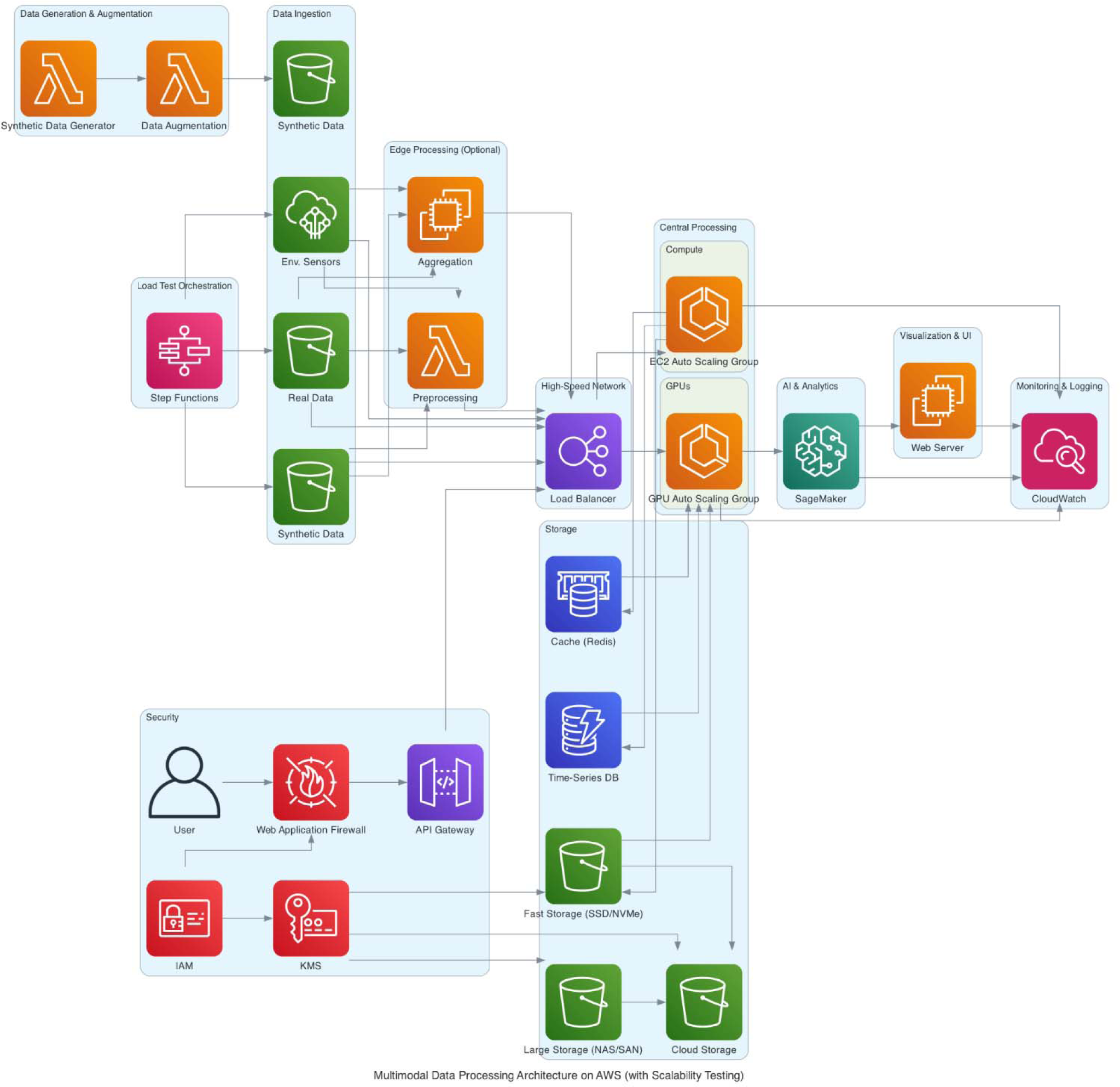
AWS Architecture depicting workflow of the model.

The overall framework scalability of the AWS services used is shown in **Figure 3b**. Thi architecture includes Amazon S3 (Simple Storage Service) that is highly scalable by default and capable of storing virtually unlimited amounts of data. The automatic scaling feature allows S3 to automatically handle increased request rates and huge volumes of data without any manual intervention.

**3b).**
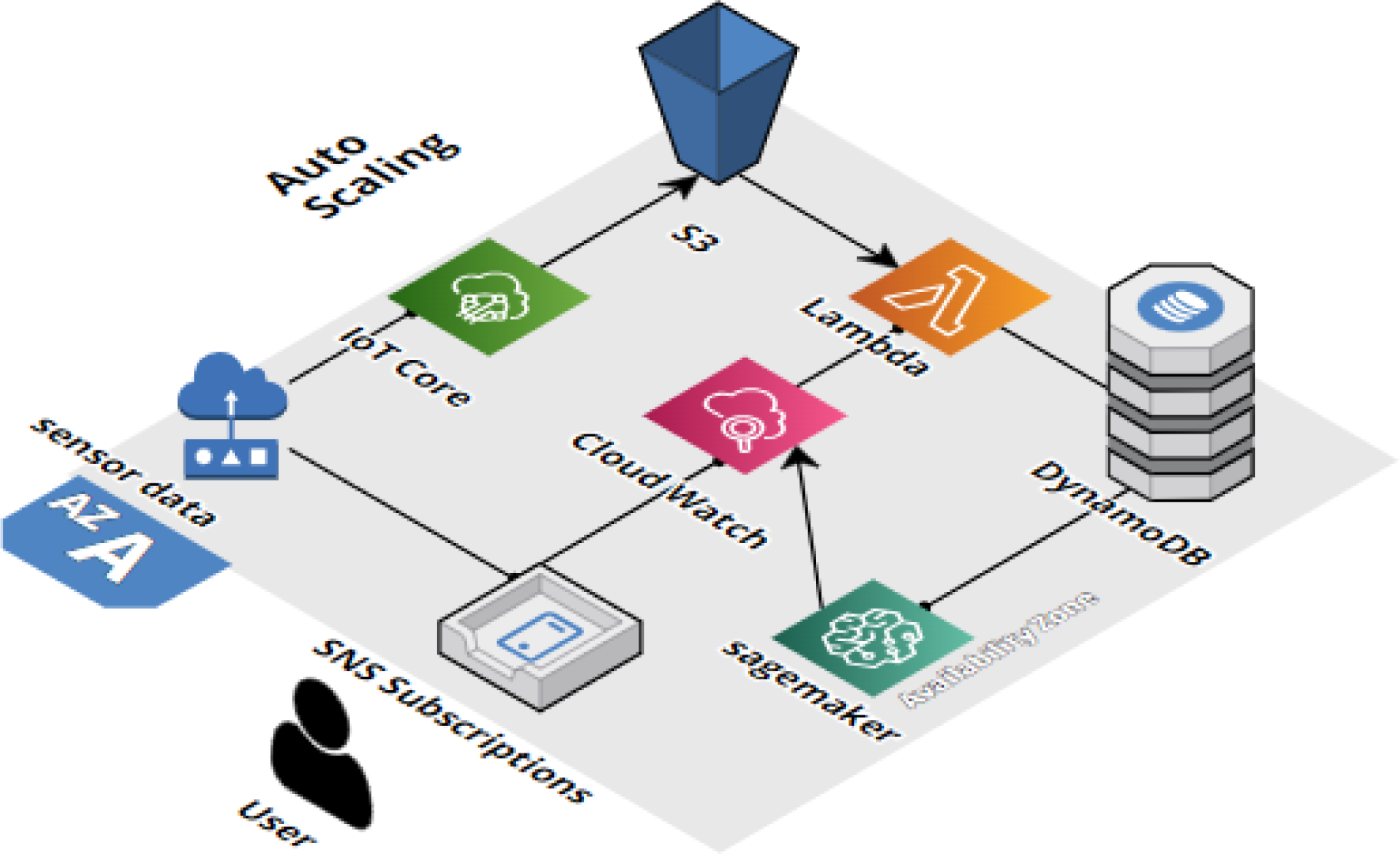
Scalability architecture of the AWS Services

DynamoDB is a fully managed NoSQL database service that can handle massive scale and high throughput. It can automatically scale its read and write capacity to accommodate changes in traffic without performance degradation. Amazon Sage Maker is a fully managed machine learning service that allows you to build, train, and deploy machine learning models at scale. It can scale horizontally to accommodate training and inference workloads across multiple instances. AWS Lambda is a serverless compute service that automatically scales to handle incoming requests and can scale horizontally to handle increased concurrency as traffic grows.

It’s pay-per-use pricing model, allows users to scale efficiently based on actual usage without overprovisioning resources.

#### 3.1.1 IoT Core Frame

The AWS IoT core consists of five services that maintain the needs of all IoT devices, connect to AWS cloud, manage devices, update over-the-air (OTA), and secure the IoT devices. It uses the TLS communication protocol to encrypt all communication. Services in this framework are rules, topics, shadow service, AWS IoT device defender, and AWS IoT device management. The Rules (**Figure 3c**) enable the IoT devices developed for smart livestock to interact with AWS services. Some of the rules used in the system are:

i. Filter incoming data from IoT devices.
ii. Simulating the IoT sensor (**Figure 3d**)
iii. Separation and recording of data according to their type in various kinds of databases.
iv. Sending notifications to users in certain circumstances (for example, the occurrence of abnormal events in the monitoring process)
v. Real-time processing of messages coming from multiple IoT devices.
vi. Setting alarms to notify the user when reaching predefined limits of certain parameters (for example, reaching a critical battery level of IoT devices, increase in HR of the animal **Figure 3e**)
vii. Send the data from a Message Queuing Telemetry Transport (MQTT) message to a machine learning frame to make predictions based on the ML model.
viii. Send data to a dashboard.

**Figure 3c).**
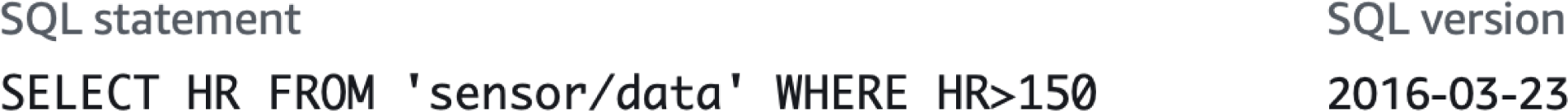
IoT Core Rule to trigger SNS if HR value exceeds a set threshold of 150 from the topic

**Figure 3d).**
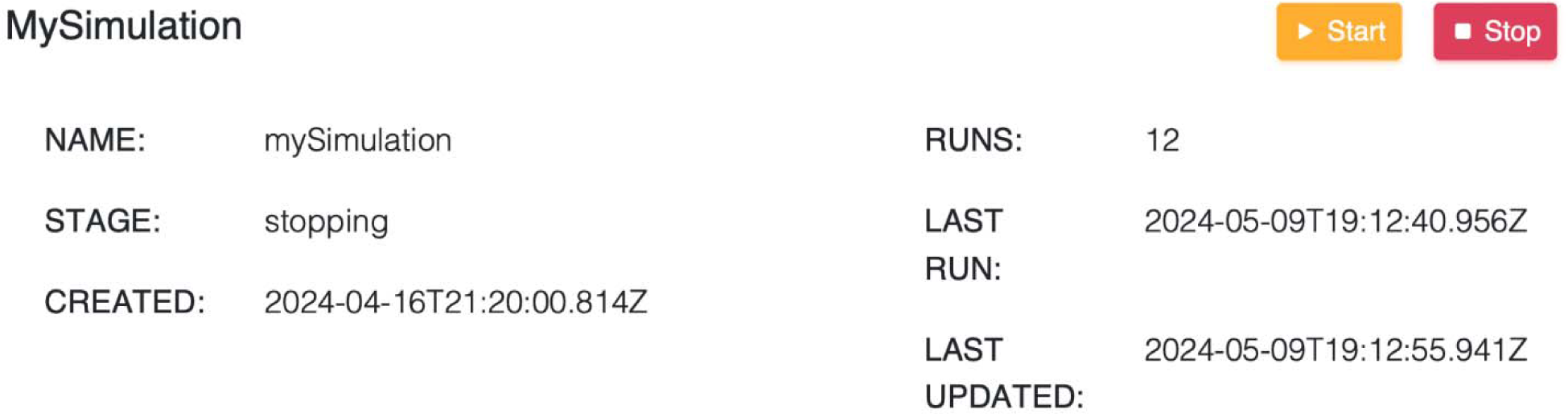
Simulation of a realtime IoT sensor to generate random values corresponding to the feature

**Figure 3e).**
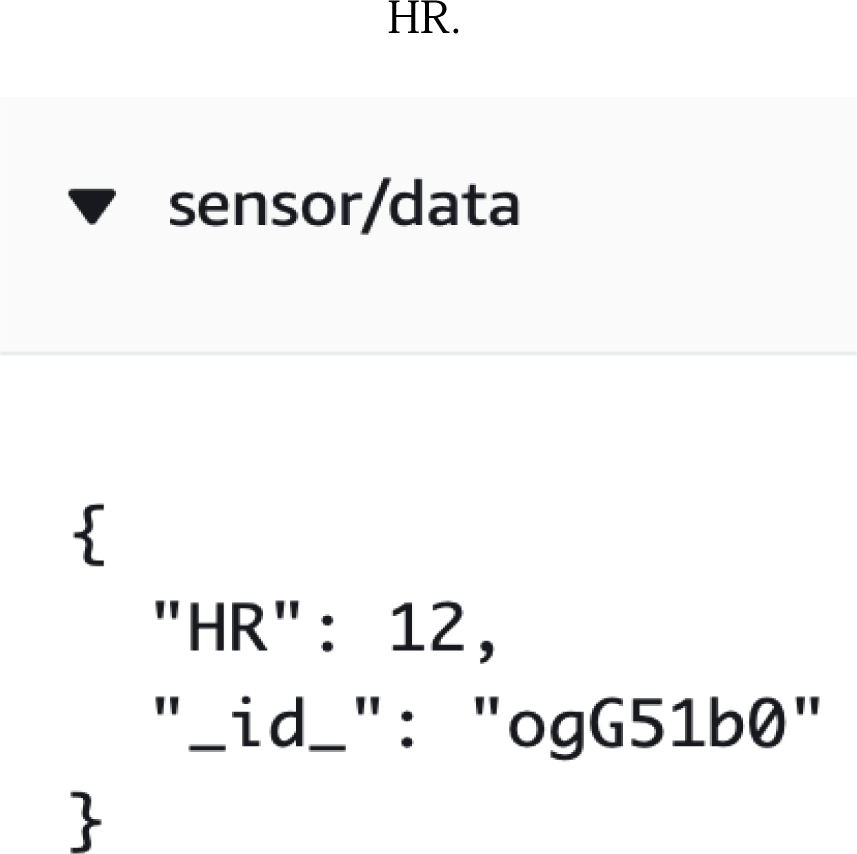
Alarm notification for the data filtered by the defined IoT Core Rule of HR greater than 150.

#### 3.1.2 Lambda Frame

The Lambda function operates as a stateless code snippet, triggered by various sources both internal and external to AWS. It offers the capability to automatically scale applications without the need for capacity planning. Unlike an EC2 instance, a Lambda serves a singular purpose and runs for a short duration, typically a few minutes. It can scale rapidly to handle hundreds of instances with minimal platform maintenance. In the context of the livestock monitoring system described in this work, Lambda functions undertake the following tasks:

Storing specific metadata, such as a unique ID, S3 bucket and key where the frame is saved, the approximate recording time, etc., to Amazon DynamoDB. Simulating IoT sensor behavior by adjusting random values to a specified frequency.

#### 3.1.3 Data Storage

DynamoDB (**Figure 3f**) is a serverless architecture are used to store events. Data from here is sent to real-time operational dashboard to provide insights. DynamoDB, with its high throughput and auto-scaling capabilities, efficiently handles large volumes of data from numerous IoT devices, ensuring the system remains responsive [73,74]. Its NoSQL design is ideal for diverse sensor data, and its high availability and durability provide robust data management. Seamless integration with other AWS services facilitates a cohesive IoT monitoring system, while low-latency operations ensure timely health interventions. Amazon Pinpoint enables effective communication and engagement through multichannel messaging, critical for real-time alerts and personalized notifications based on specific data points [75]. Integration with AWS IoT Core and Lambda triggers real-time alerts for anomalies, supporting automated health checks and scheduled notifications. Detailed analytics on message delivery and user engagement refine communication strategies, ensuring critical alerts are promptly addressed. Pinpoint’s scalability accommodates extensive livestock operations, offering targeted messaging for efficient health management. Together, DynamoDB and Amazon Pinpoint provide a robust, scalable, and efficient system for livestock health monitoring, enhancing animal welfare and farm productivity through effective data management and real-time communication.

**Figure 3f).**
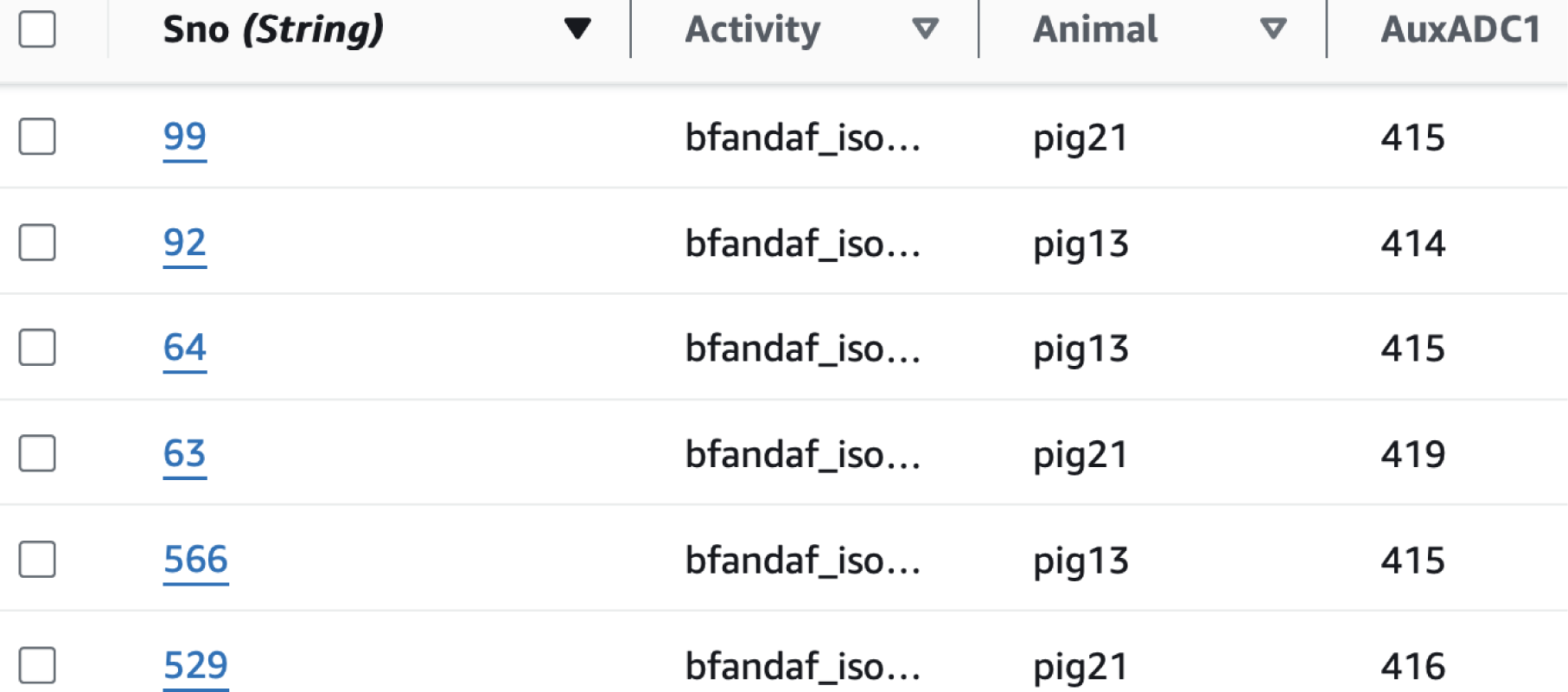
Dynamodb creates tables that can store and retrieve any amount of data and serve any level of request traffic.

#### 3.1.4 Notification Frame

Amazon Pinpoint, as a communication service, connects with users through various channels such as email, SMS, push, or voice (**Figure 3g**). This service is used to personalise messages with the right content [76]. It sends push notifications to the smart livestock application after pre-provided data that authorises PinPoint to send messages. The credentials that are provided depend on the operating system. For iOS apps, an SSL certificate is provided. The certificate authorises the PinPoint service for sending messages to the smart livestock apps. For Android apps, a web API key is provided. These credentials authorise the PinPoint service for sending messages to the smart livestock apps. AWS AppSync takes care of managing and updating real-time data between web and mobile app and cloud. Additionally, it allows apps to interact with data on mobile devices when it is offline.

**Figure 3g).**
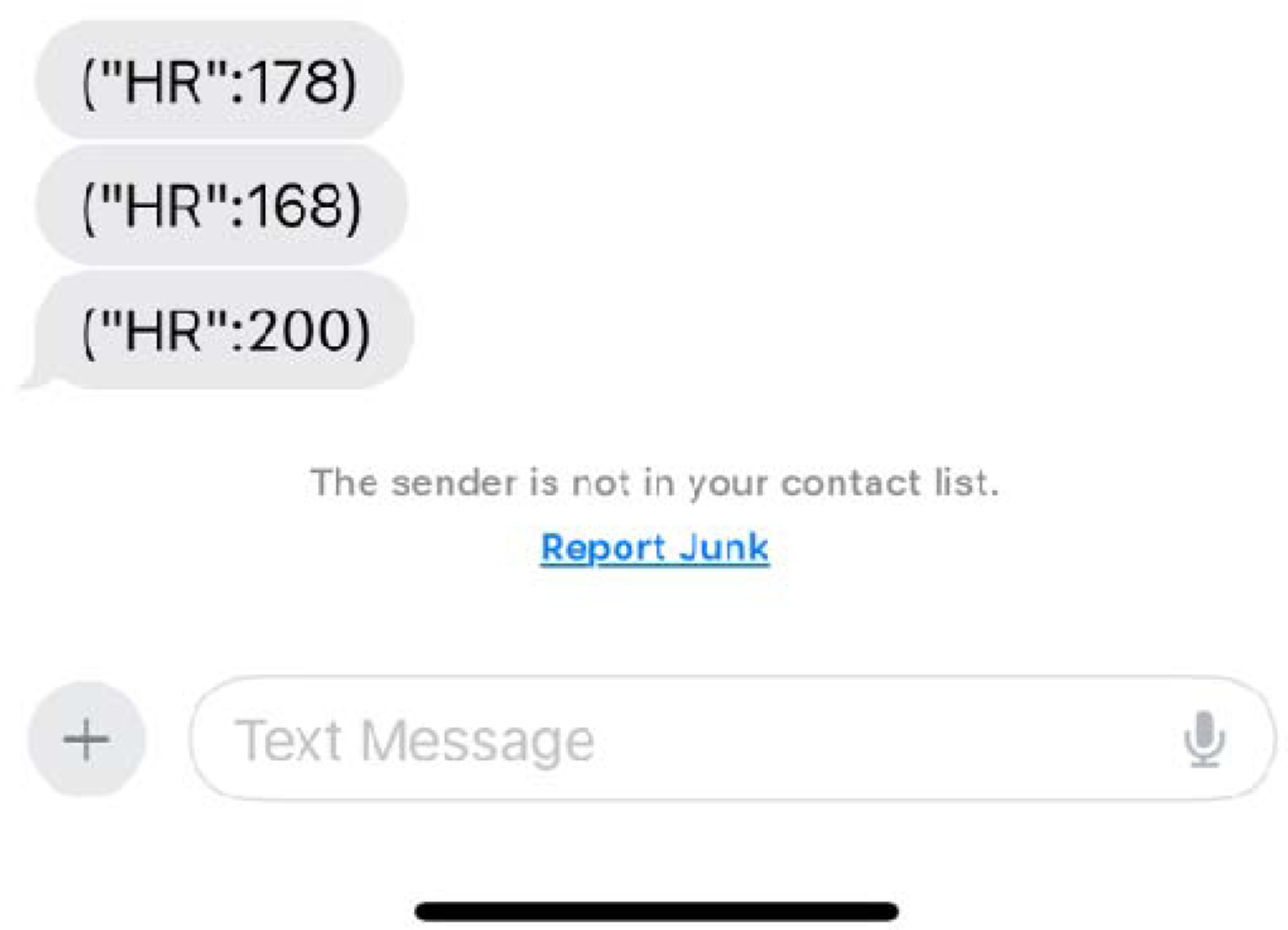
A screenshot representing IoT rule sending an alert message to the user as the HR values exceeding the set threshold of 150 [77].

#### 3.1.5 Machine Learning Frame

Long-term unused data can be stored in S3 Glacier which is designed to store rarely used data.

Since the data from S3 Glacier is not readily available it is archived. The unused data can be unzipped and moved to S3 bucket if required. The data from S3 is then used by Amazon Sage Maker to build, train, and deploy interface models [78]. To predict the future heart rate of the animals, the model is trained, which takes as parameters the animal’s temperature, heart rate, isolation, and pairing.In addition to the classification of the health status of the animals, a regression is performed to predict, in the short-term, the future amount of milk that will be extracted from each animal.

### 3.2 Execution Results

#### 3.2.1 IoT Device

A group of sensor groups, logical blocks, and power supply collected sensor readings from livestock for temperature, humidity and heart rate. The communication in each livestock IoT device is device-to-edge using Nb IoT [79]. All IoT devices on a farm have strict firewall rules. The various sensor devices deployed are registered prior to initialization of the framework for security. Livestock IoT devices will not be able to communicate with each other on their farm or with other devices to ensure independent working of each sensor. In case the IoT device cannot connect to the IoT edge, or the data transmission is obstructed by any other reasons, the device can keep the sensor readings until the moment it reconnects. Then, the IoT device will send the up-to-date data.

#### 3.2.2 IoT Edge

The AWS IoT Core, which is an Internet of Things (IoT) open-source edge runtime and cloud service that helps to build, deploy, and manage device software [80] . It was used to manage local processes, communicate, and synchronise certain groups of devices and exchange tokens between Edge and cloud, which acts as a hub or gateway in Edge. The communication is Edge-to-Cloud using TCP based protocols. It consists of MQTT Broker, Local Shadow Service, AWS Lambda, Meta Data, and Trained Models. AS IoT Core was used to perform the following tasks:

i. Processing of large data streams and automatically sending them to the cloud through Lambda functions (AWS Lambda);
ii. Lambda functions uses Nb IoT protocol for connections between livestock IoT devices and cloud using device authentication and authorisation (Meta Data);
iii. Deployment of cloud-trained machine learning models for regression that predicts a percentage of the future power of the battery concerning the individual frequency and load of the monitoring livestock system (Trained Model);
iv. Updated group configuration with secured over-the-air (OTA) software updates.

The developed framework is tested against three different sizes of data; The dataset contains 318713*44, 320713*44, and 322713*44 in scenarios 1,2 and 3 respectively.

The execution results from Lambda displaying a) Error count and success rate(%); b)Throttles; c)Invocations; d)Duration; e)Total concurrent executions; in three different scenarios of changing farm size are as follows.

It can be inferred from the Lambda metrics that a high success rate suggests that Lambda functions are functioning as expected and handling incoming requests effectively with increasing data load **Figure 3.6h**.

Throttle lines **Figure 3.6i** intersecting or crossing the x-axis indicate instances when Lambd invocations were throttled. The position of these lines along the x-axis corresponds to th timestamp or time period when throttling occurred.

**3.6h) (i).**
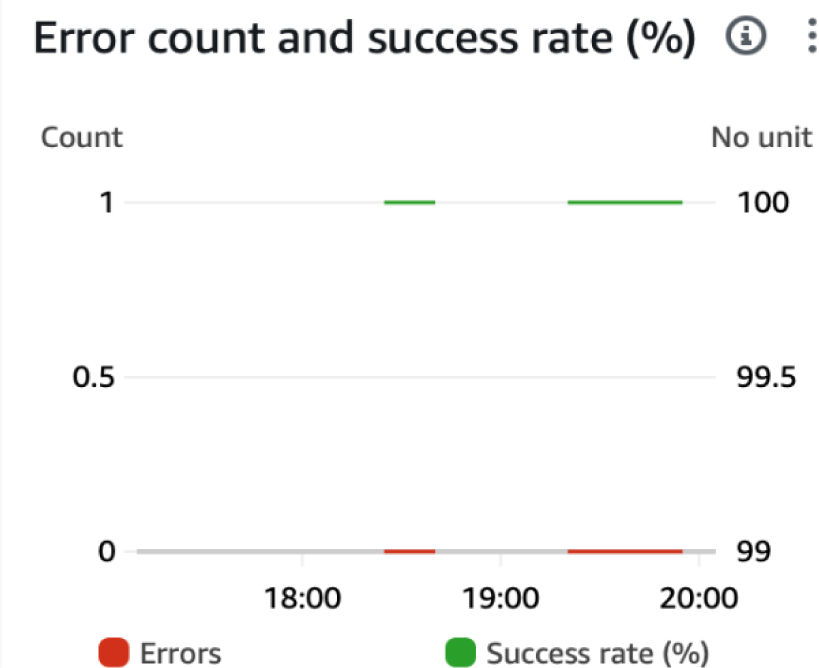
Lambda execution results displaying the error and success rates when experiencing data load in scenario 1.

**3.6h) (ii).**
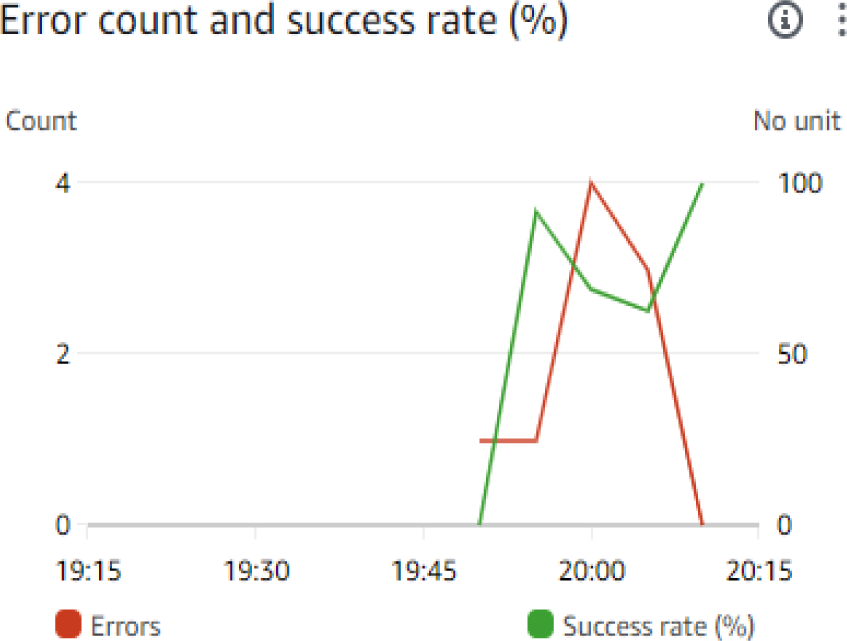
Lambda execution results displaying the error and success rates when experiencing data load in scenario 2.

**3.6h) (iii).**
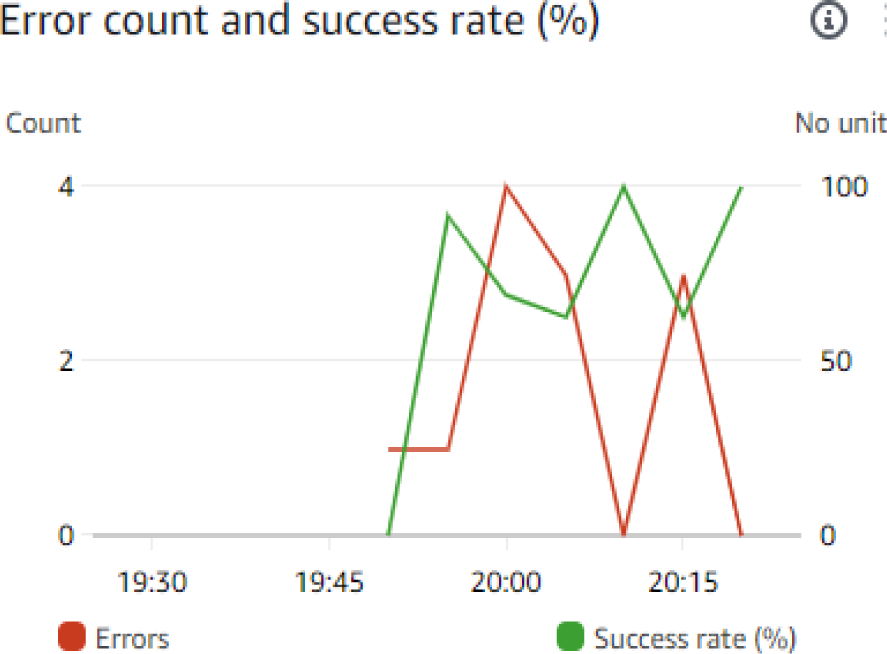
Lambda execution results displaying the error and success rates when experiencing data load in scenario 3.

**Figure 3.6i) (i).**
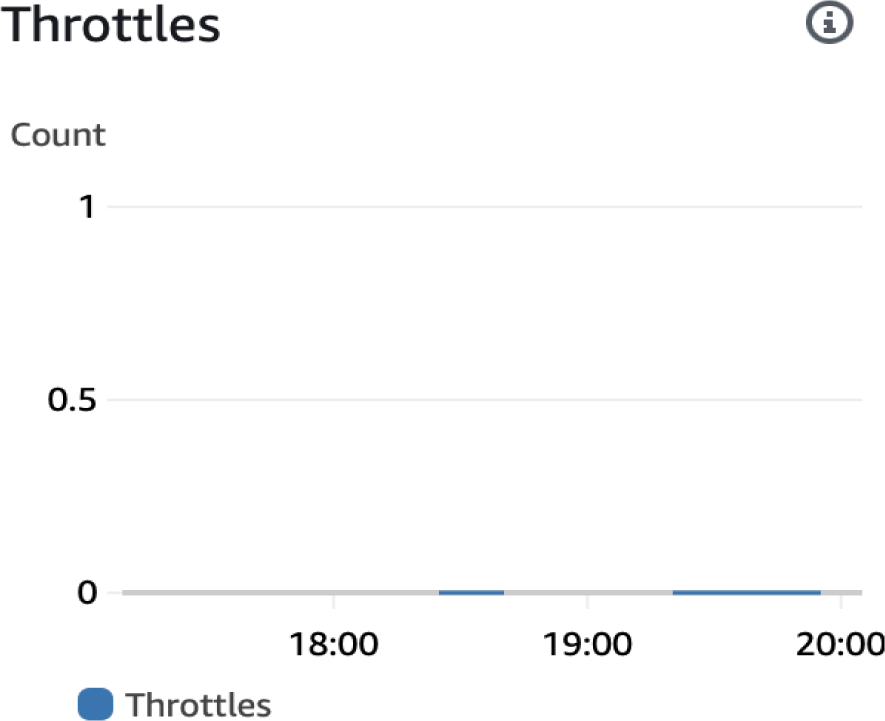
The count of throttles in Lambda function with small data load.

**Figure 3.6i) (iii).**
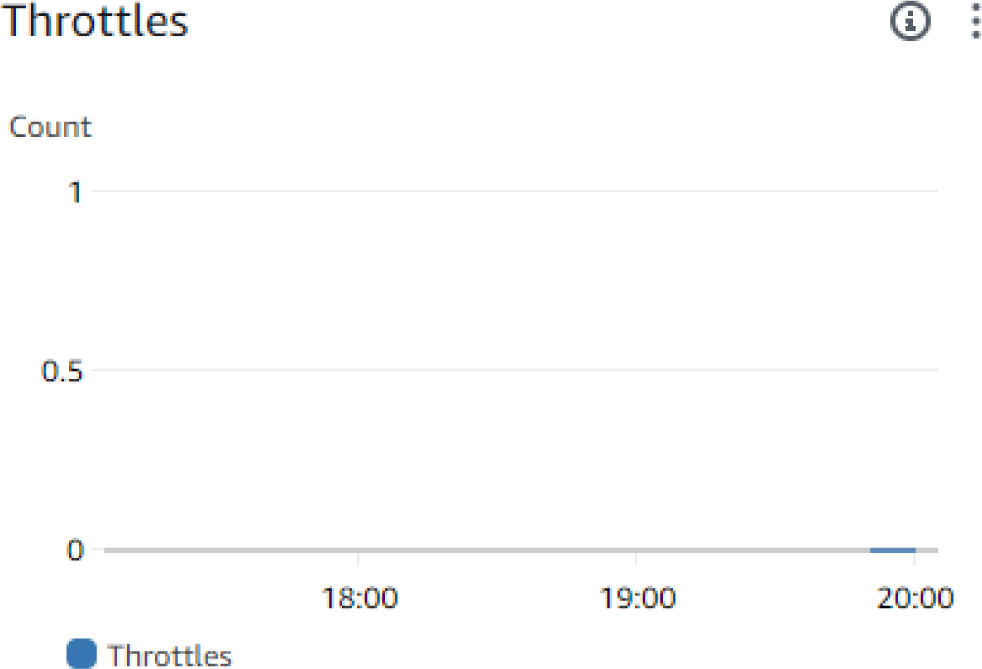
The count of throttles in Lambda function with medium data load.

**Figure 3.6i) (iii).**
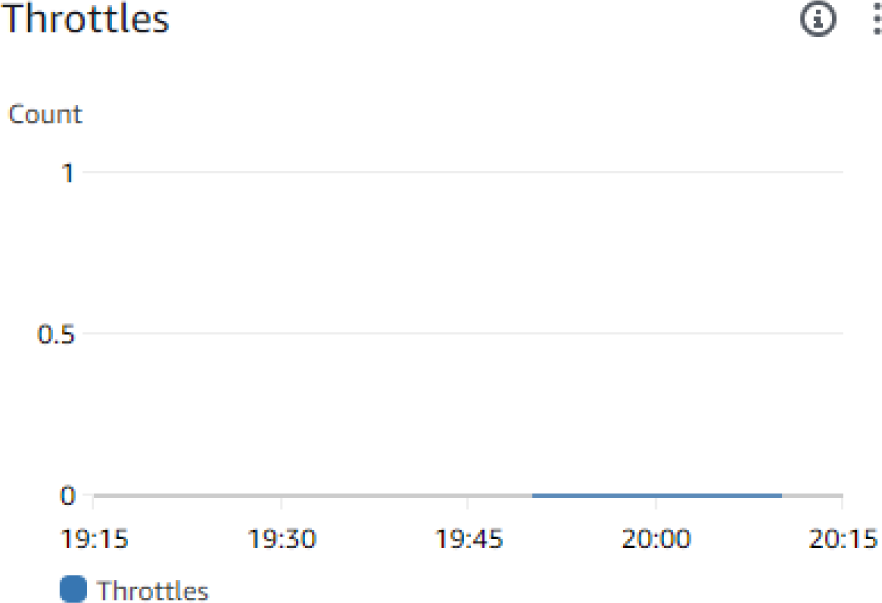
The count of throttles in Lambda function with large data load.

The above screenshot depicts that the Lambda function is operating efficiently and effectively, with no restrictions on the number of concurrent executions it can handle. It is also evident that the application can horizontally scale effectively to accommodate varying workloads without being hindered by concurrency limitations. AWS Lambda is utilizing the available resources (such as memory, CPU, and networking) to their fullest extent, maximizing the efficiency of your function executions.

**Figure 3.6j** shows Lambda function is triggered each time a service is triggered. Here, whenever S3 gets a new data added, Lambda function is invoked and automatically scales to accommodate incoming requests. As the number of invocations increases, AWS Lambda manages the allocation of resources to handle the workload efficiently.

**Figure 3.6j) (i).**
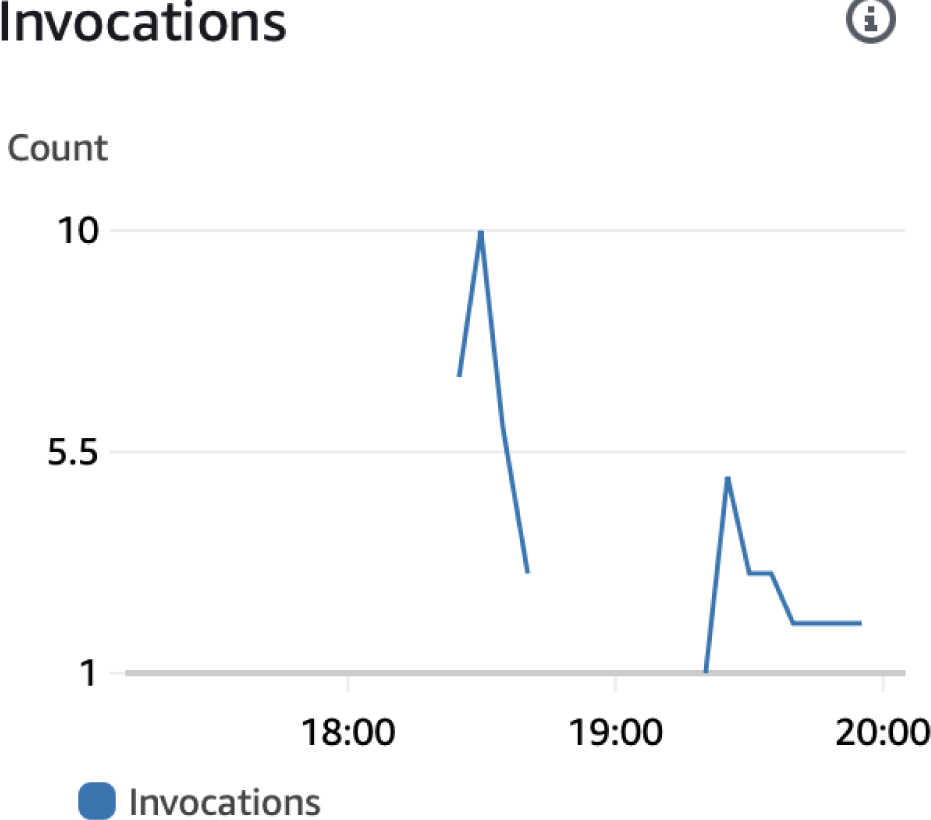
Examining Lambda invocations in Light data load.

**Figure 3.6j) (ii).**
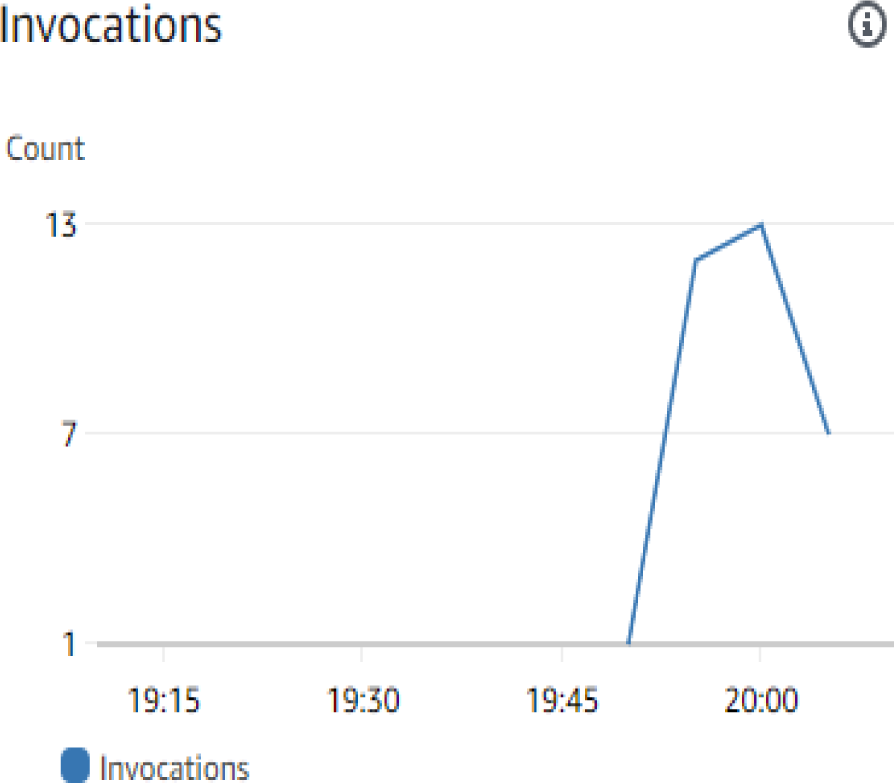
Examining Lambda invocations in moderate data load.

**Figure 3.6j) (iii).**
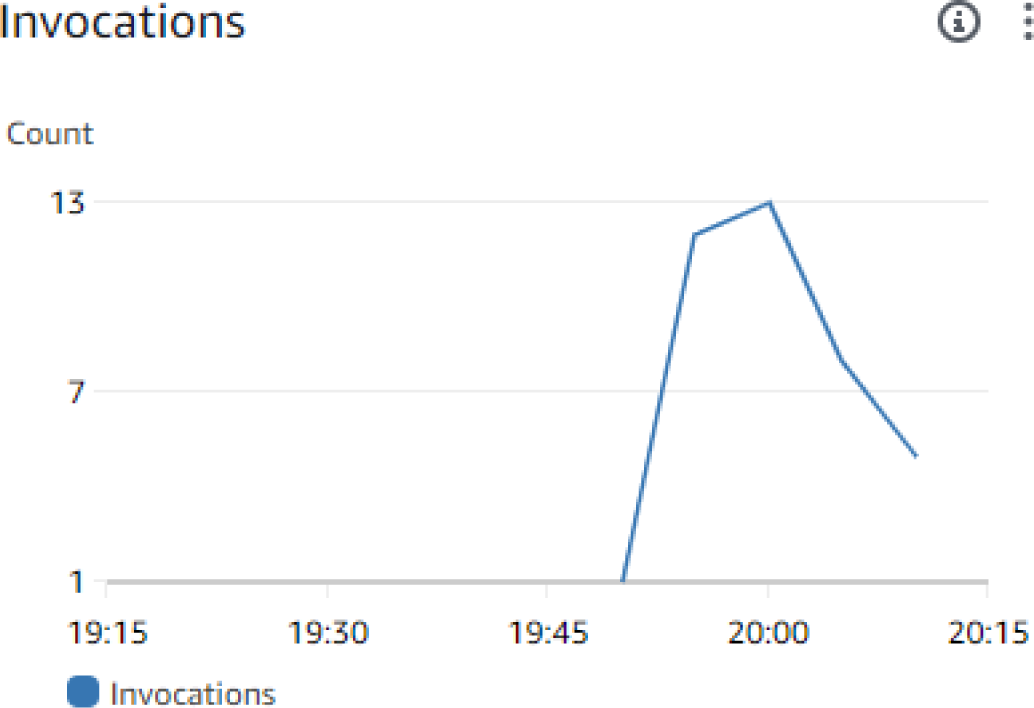
Examining Lambda invocations in heavy data load

In AWS Lambda, “duration” refers to the time taken for a Lambda function to execute a single invocation from start to finish **Figure 3.6k**. It includes the time it takes to initialize the execution environment, execute the function’s code, and handle any outgoing requests or asynchronous operations.

**3.6k) (i).**
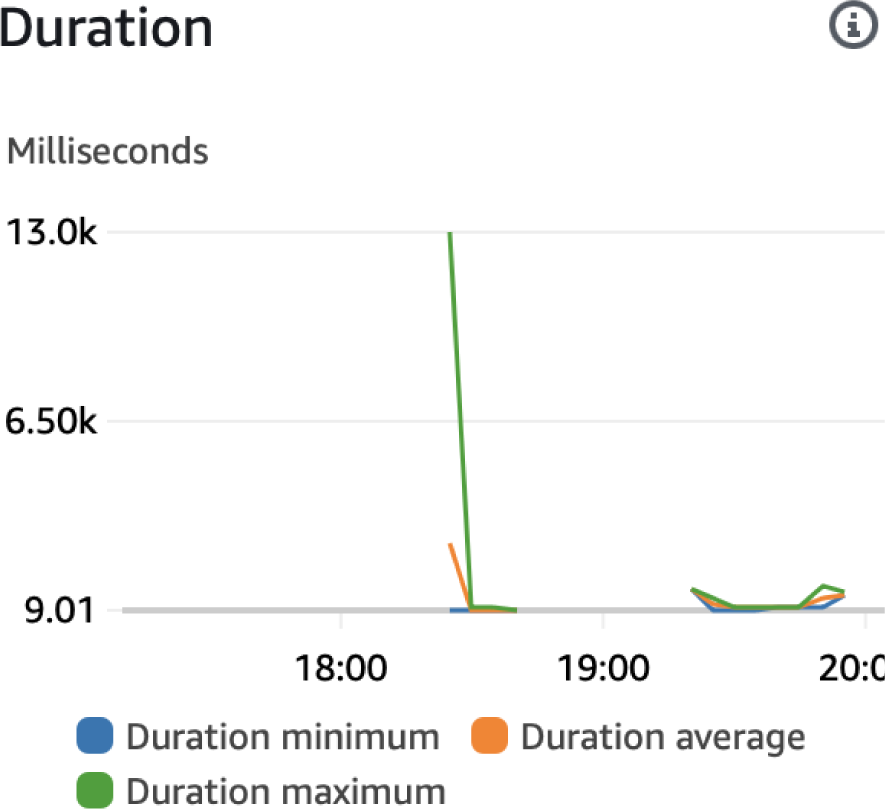
Duration variability in case 1.

**3.6k) (ii).**
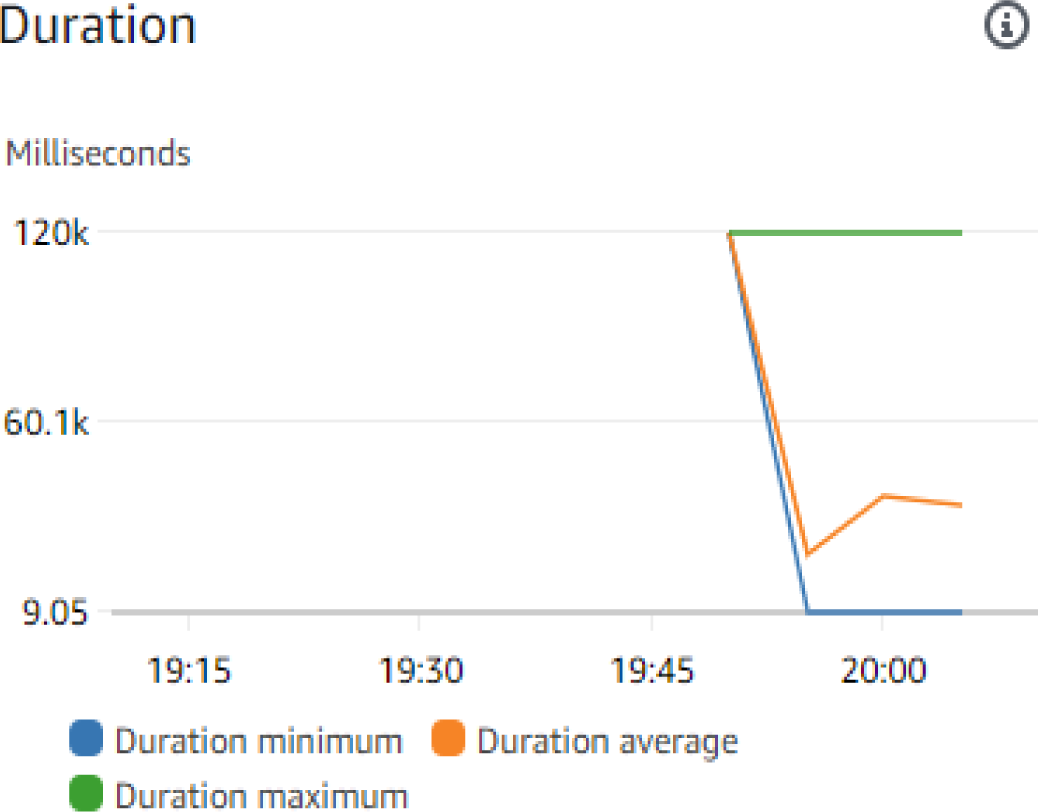
Duration variability in case 2.

**Figure 3.6k) (iii).**
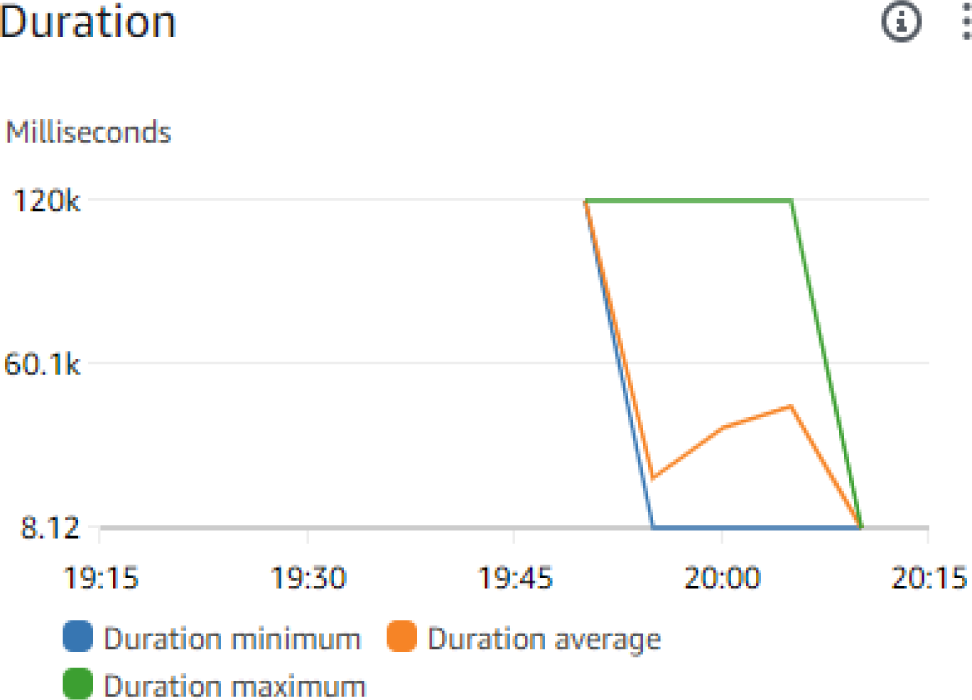
Duration variability in case 3.

Optimizing code, memory allocation, and concurrency settings can lead to improved performance, reduced costs, and enhanced user experiences in serverless applications **Figure 3.6l** depicts the total concurrent executions that occurred. Peaks in the graph represent times when the total concurrent executions reach their highest levels. This occurs during periods of high traffic, increased workload, or bursts of activity Built on the AWS cloud platform, which provides more than 200 services, is the smart livestock architecture. More than 20 management services, over 30 machine learning and data analytics services, and over 13 database and storage services are included in this list. This research’ recommended architecture uses AWS serverless services to make it easier to create vital data pipelines that can effectively handle massive amounts of data from IoT devices. It also makes it possible to store large amounts of unprocessed raw sensor and image data on AWS S3 at a reasonable price while maintaining accessibility to both old and new data. **Figure 3.6m** displays the error and success rate recorded in Cloud Watch which is used to analyze the performance and reliability of the model.

**3.6l) (i).**
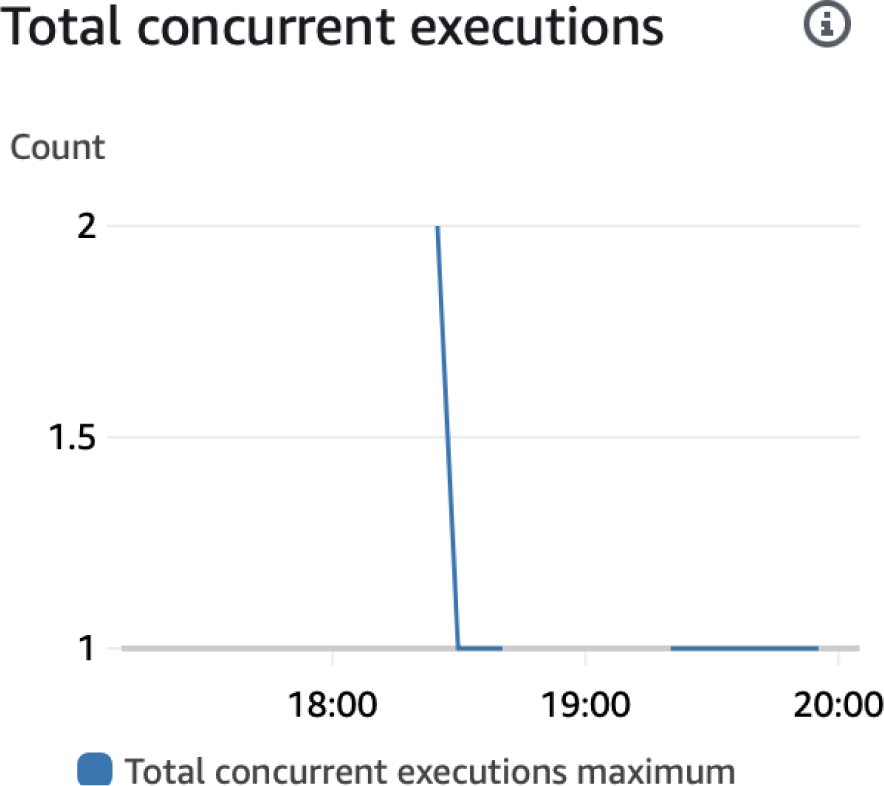
Total Concurrent Executions with with Small Data Size.

**3.6l) (ii).**
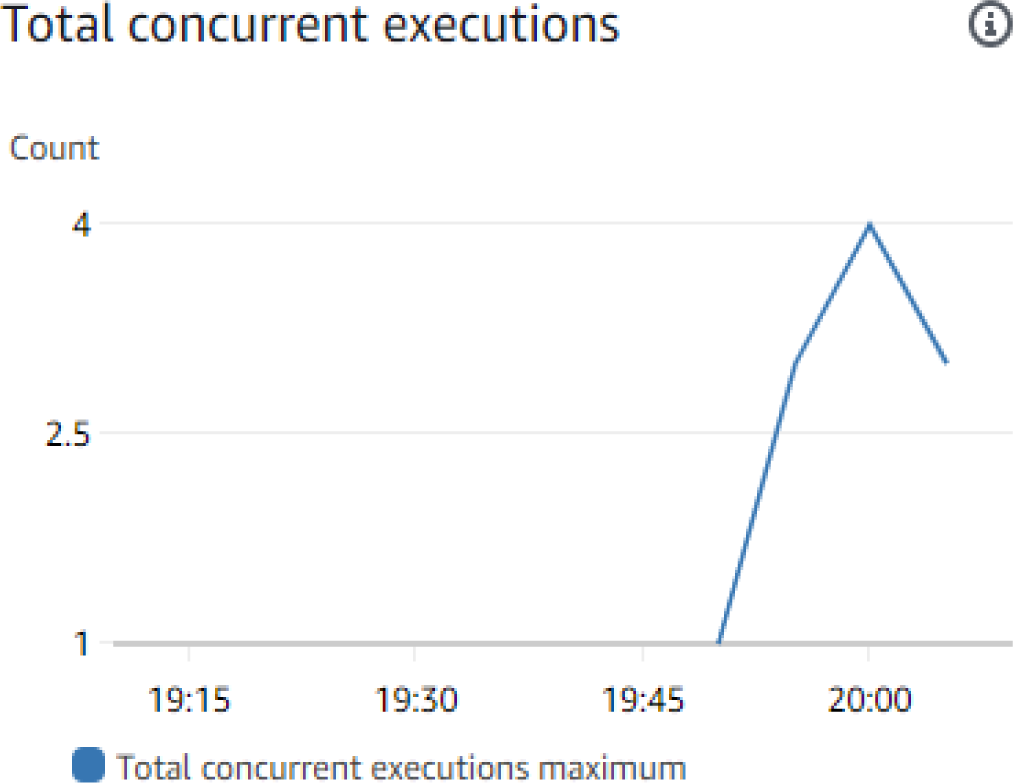
Total Concurrent Executions with Medium Data Size.

**3.6l) (iii).**
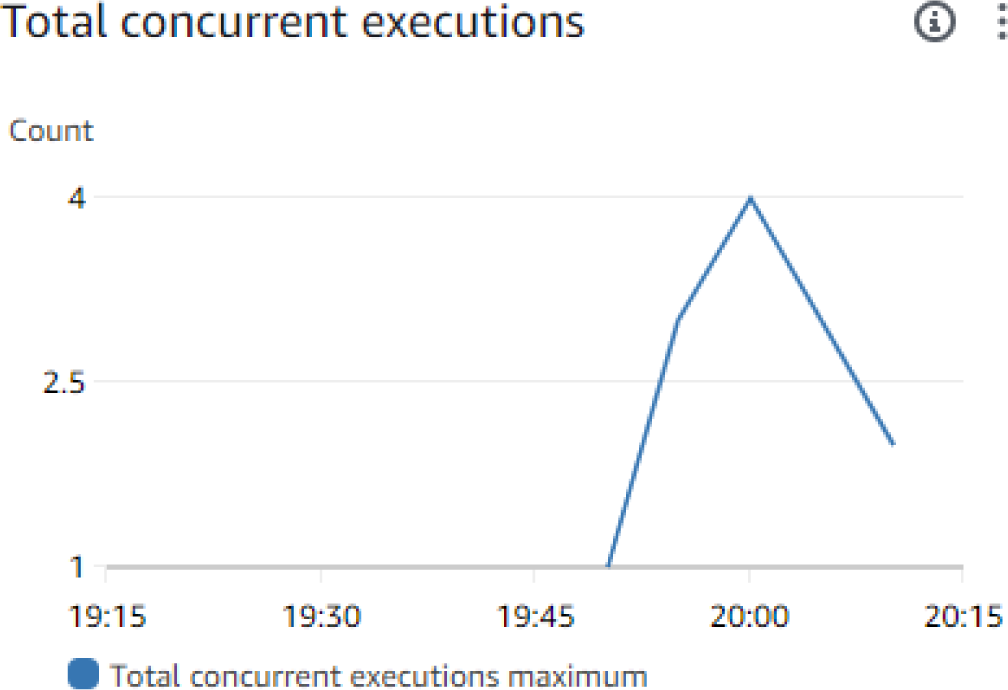
Total Concurrent Executions with Large Data Size.

**Figure 3.6m) (i).**
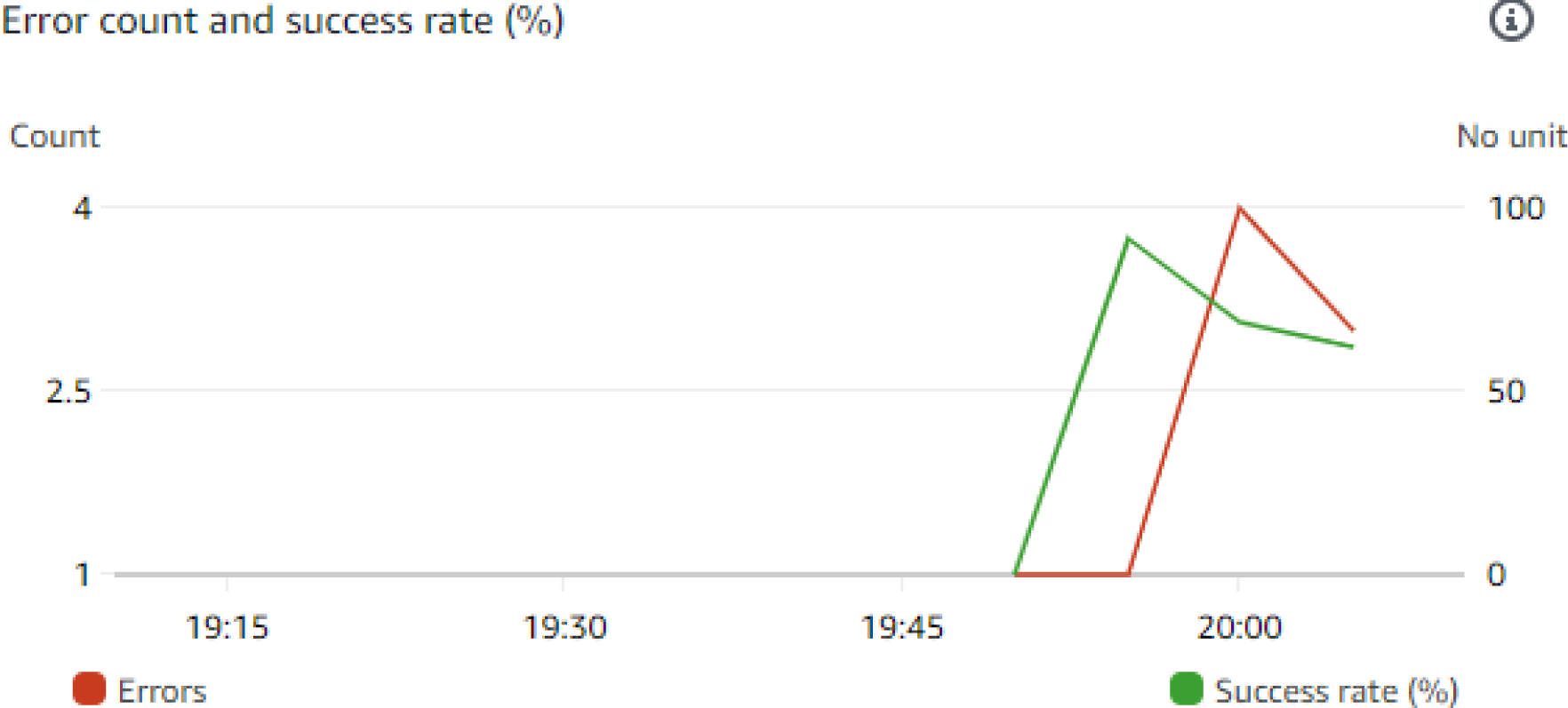
Cloud Watch metric with small data load.

**Figure 3.6m).**
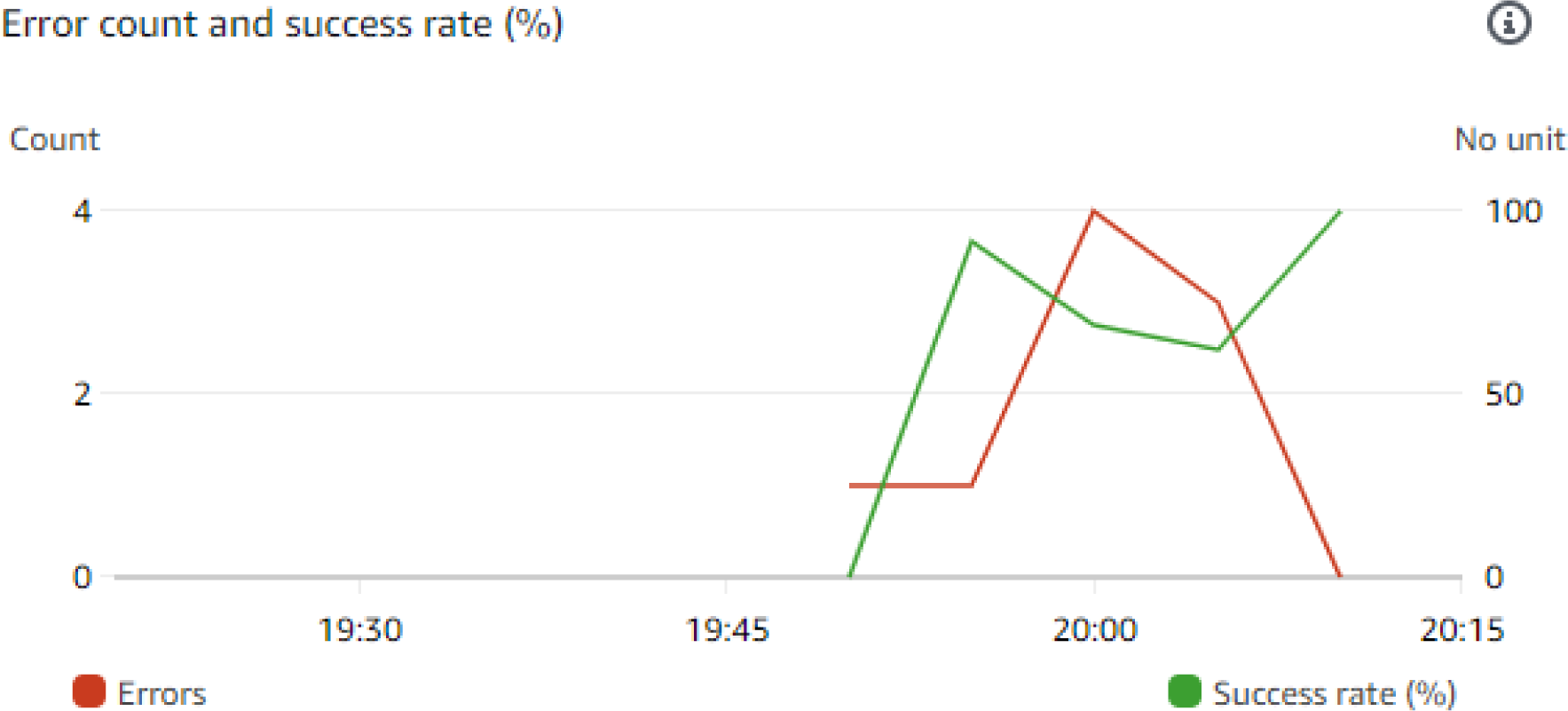
Cloud Watch metric with large data load.

The developed model predicts the heart rate considering various parameters including animal activity, temperature, breathing rate etc. **Figure 3.6n** shows the predicted and actual values of the HR using Sage Maker.

**3.6n).**
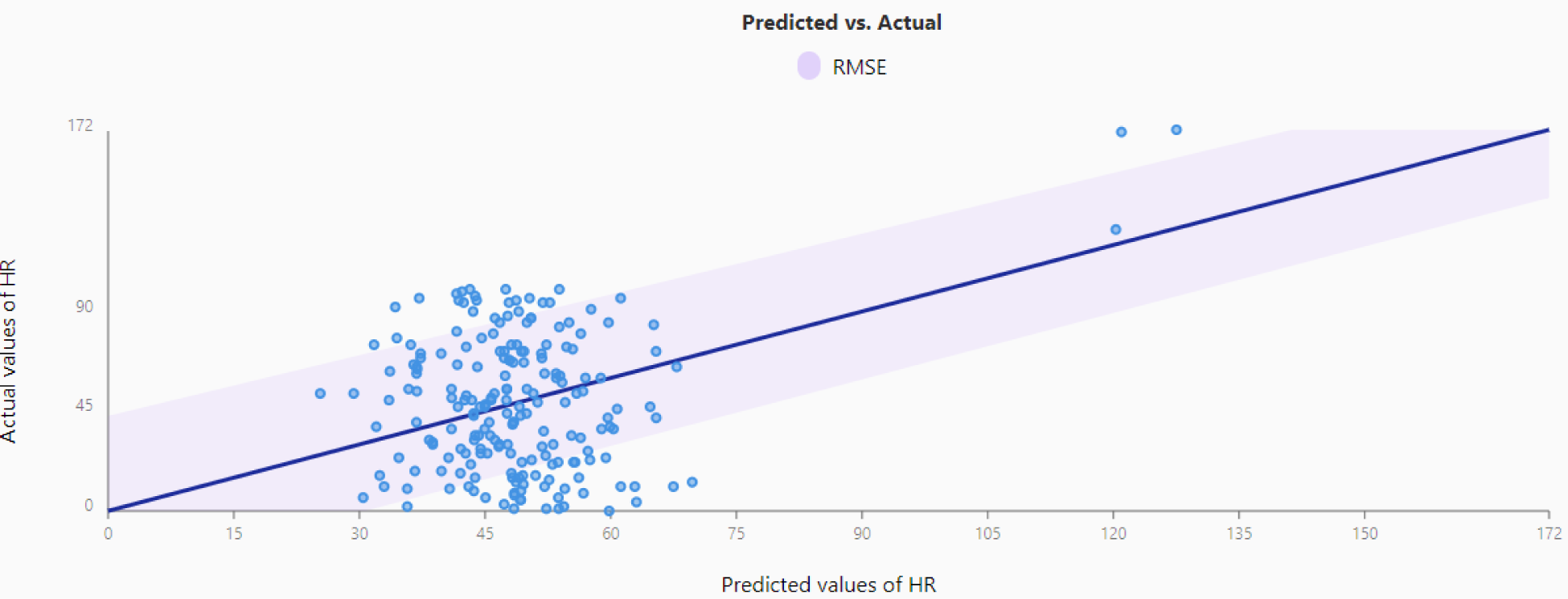
Graph displaying RMSE of predicted and actual values of HR.

Among all the features available to predict HR, the significant contributors are Breathing Waveform Figure 3.6o and Activity Figure 3.6p .

**3.6o).**
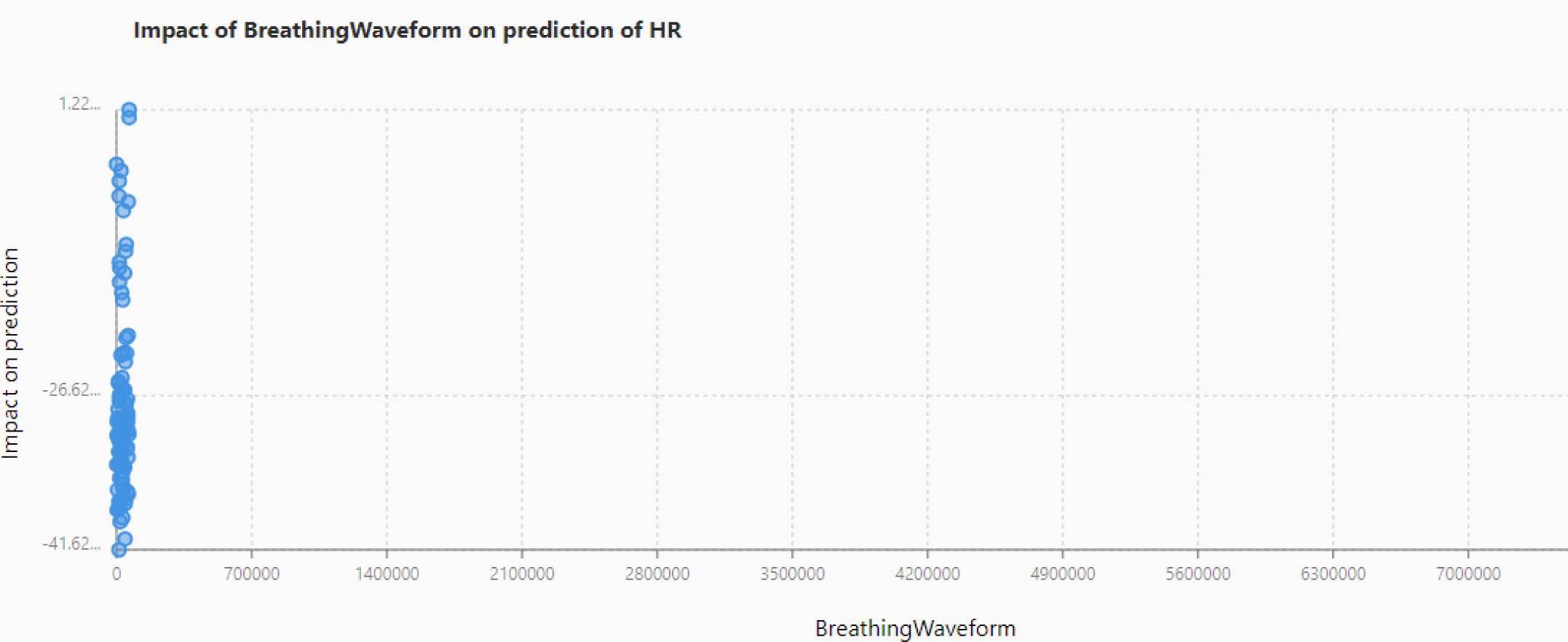
The plotted graph shows the Breathing Waveform in the x-axis and its impact on prediction on the y-axis in forecasting HR value.

**3.6p).**
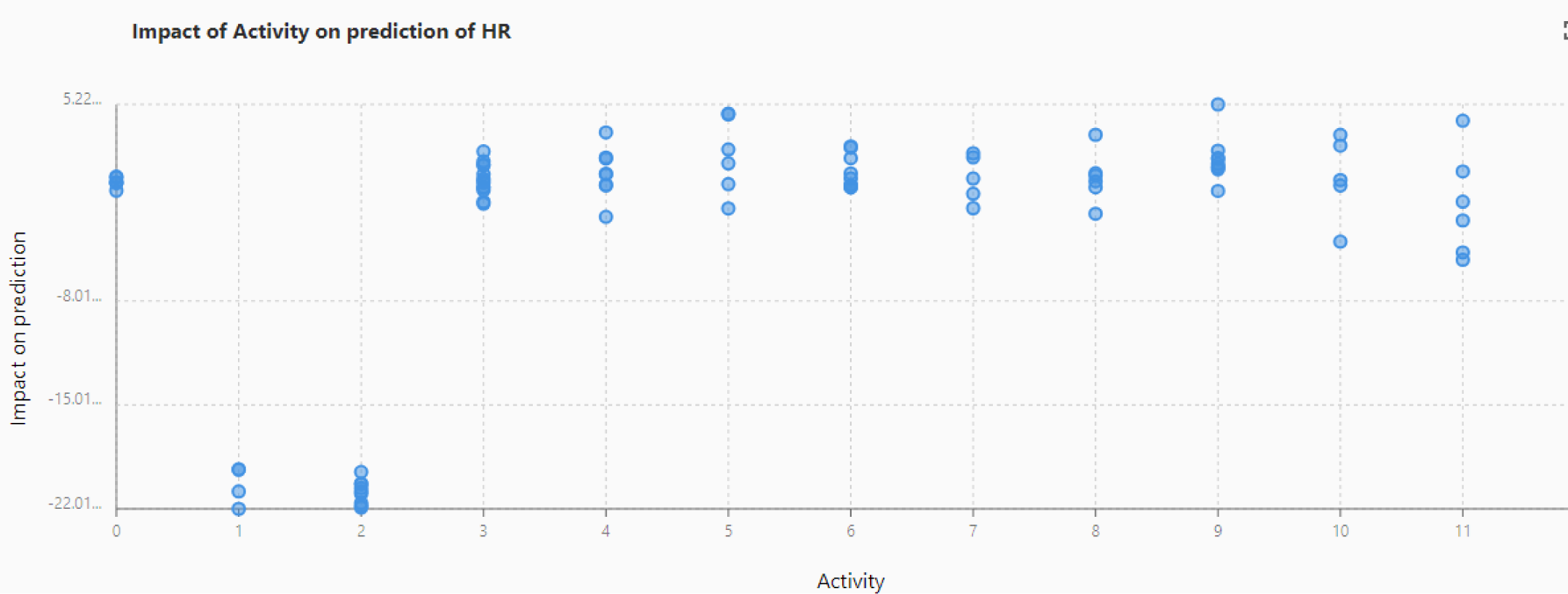
The graph displays the Activity of the animal against the impact of prediction to predict the values of HR.

## 4. Future Work

The landscape of Internet of Things (IoT) systems is marked by a rich array of sensors and communication protocols, which collectively form the backbone of highly efficient ecosystems.

Central to this technological frontier is the concept of digital twin technology, designed to create virtual replicas of physical systems for in-depth analysis and optimization. This approach provides a profound opportunity to enhance system monitoring and management across various sectors, including agriculture. Focusing on specific technologies, LoRaWAN and NB-IoT stand out for their unique capabilities and integration potential. LoRaWAN is particularly adept at enabling long-range communication with minimal power consumption, making it ideal for monitoring devices across extensive areas such as agricultural fields, where energy efficiency is paramount. This technology forms a foundational element of digital twin architectures, facilitating the remote observation and management of devices in energy-limited scenarios. Conversely, while NB-IoT may present higher initial costs and complexity than LoRaWAN, its superior coverage and energy efficiency provide significant benefits. In the context of digital twins, the extensive data required to accurately reflect complex systems necessitates robust and reliable communication technologies. NB-IoT, supported by major telecommunication networks, offers high fidelity in asset monitoring and enables consistent data collection from a diverse array of sensors. This capability is critical for real-time analysis and the effective implementation of predictive maintenance strategies. Ensuring the optimization of the architecture when excessive data load reaches the model is unanswered. This shall ensure performance optimization and effective cost-cutting. Looking to the future, the convergence of IoT with digital twin technology holds the promise of transforming predictive capabilities in numerous domains. By integrating with advanced artificial intelligence (AI) and machine learning algorithms, these technologies could foster the development of self-optimizing systems. Such systems would not only replicate current conditions but also anticipate and adapt to future.

## 5. Conclusions

In the realm of agricultural technology, automated smart livestock monitoring systems represent a frontier yet to be fully conquered. While substantial technological advancements have been achieved, the seamless integration of these systems into daily farm operations with minimal human oversight remains a challenging endeavor. This study has taken significant strides toward overcoming these challenges by developing and refining an intelligent livestock monitoring architecture. Our extensive testing of the system’s most critical aspects—from sensor accuracy to network resilience—has confirmed its robust functionality and reliability. These positive outcomes demonstrate the system’s readiness for broader adoption in livestock farms. The core of our contribution through this paper is the presentation of a novel IoT architecture that stands out due to its independence from the IoT device layer. This design is not only innovative but highly adaptable, supporting a wide variety of data formats, enhancing feature extensibility, and accommodating diverse communication protocols. One of the key strengths of this architecture is its ability to adapt to future technological changes. By decoupling the IoT layer from the cloud, we have created a system that can easily integrate new devices or update communication protocols without disrupting existing operations. This flexibility is crucial for maintaining the longevity and relevance of the system as technological standards evolve. Additionally, the use of Narrowband IoT (Nb-IoT) technology is a pivotal aspect of our system design. Nb-IoT enhances the system by minimizing power consumption, which is crucial for long-term, sustainable operations in remote or difficult-to-access areas. This technology also extends the coverage area well beyond that of traditional monitoring systems, reaching deep indoor environments where conventional signals might falter. Moreover, the inherent simplicity of the Nb-IoT devices ensures that the system remains manageable and scalable, even as farm operations grow or as the technology landscape shifts. These enhancements made possible by Nb-IoT not only meet but exceed the current requirements for modern livestock management, offering unprecedented levels of operational efficiency, data reliability, and system scalability. This robust validation of the architecture marks a significant milestone in the journey towards automated, smart livestock management systems and sets a new benchmark for the industry, promising enhanced predictive analytics, and operational efficiencies. The practical implications of these advancements are profound, signaling a shift towards more proactive and precision-based farming practices that could redefine livestock management in the years to come.

## Acknowledgments

The authors gratefully acknowledge the financial support provided by the Natural Sciences and Engineering Research Council of Canada and Agriculture and Agri-Food Canada’s Sustainable Canadian Agricultural Partnership program. We also extend our sincere thanks to Professor Israat Haque from the Faculty of Computer Science at Dalhousie University for her valuable insights, critical remarks, and thorough review of our manuscript, which significantly improved its readability.

## Appendix Supplementary materials

**Table 1**.

## Data availability

Data will be made available on request.

