## Supplementary material for "Development of a Cloud-Based IoT System for Livestock Health Monitoring Using AWS and Python": Table S1 Supplementary File

**Supplementary Information**

Table 1. Comprehensive Overview of IoT and Cloud Technologies in Livestock Health Monitoring

| **References** | **Objective** | **Parameters** | **Type of Sensor/Technology** |
| --- | --- | --- | --- |
| [24] | Automated dairy cow health evaluation system development based on IoT. | Ruminating, feeding, sleeping, and moving | Rumination sensor, accelerometer, Wifi |
| [25] | System demonstration for recording cow body measurements and milk yield prediction. | Body temperature, relative humidity, heart rate, rumination rate | Temperature sensor, heart rate sensor, humidity sensor, rumination sensor, Atmega328p, NodeMCU |
| [26] | IoT application in cow health, milk processing, and transport security. | Real-time location, temperature, heart rate, breath, rumination | Temperature sensor, heart rate sensor, rumination sensor, Bluetooth Low Energy (BLE), ZigBee |
| [27] | Discussion on IoT and AI-integrated cattle health monitoring devices. | Body temperature, heart rate, rumination | Heart rate sensor, temperature sensor, rumination sensor, ZigBee, WSN |
| [28] | Introduction of the “MOOnitor”, an intelligent IoT device for cow health monitoring. | Estrus monitoring, temperature, diseases | Temperature sensor, GPS, accelerometer, GSM module |
| [29] | Discussion on IoT for modern swine health farming. | Feed intake, temperature, cough, body temperature | RFID, Zigbee, GPS |
| [30] | Evidence-based approach to the monitoring and surveillance of farmed animal welfare. | Identification of pigs, individual animal data, date of birth, mortality, etc. | RFID |
| [31] | Review of IoT technologies for swine farming. | Temperature of individual pigs or whole herd, muscle injuries, infectious diseases, ovulation | Infrared thermal imaging |
| [32] | Real-time monitoring of environmental parameters in a gestating sow house. | Temperature, relative Humidity, CO2 and ammonia concentrations | Zigbee |
| [33] | Automated pig feeding and behaviour tracking. | Auto locomotion, movement pattern, behavior, posture, tail biting, etc. | Deep learning/image analysis |
| [34] | Review of accelerometer for cattle and pig welfare. | Pigs’ movement pattern, standing time, posture, etc. | Accelerometer |
| [35] | Web app for smart agriculture using LoRa and LoRaWAN. | Monitor and regulate humidity and temperature levels, automating temperature and irrigation. | PLC, Cloud, LoRaWAN |
| [36] | Cattle Monitoring using LoRaWAN. | Monitor the movement of cattle. | IoT, GPS, LoRaWAN, Cloud |
| [37] | Using LoRaWAN for Livestock Monitoring with drones. | Aerial monitoring of farm and animals | Drones, LoRaWAN |
| [38] | Health monitoring and Animal-vehicle collision using LoRaWAN. | Health monitoring | LoRaWAN, GPS, WSN |
| [39] | LHM using LoRaWAN. | Health monitoring | IoT, LoRaWAN, GPS |
| [40] | Livestock location monitoring using LoRaWAN. | Monitor movement of livestock | IoT, LoRaWAN, GPS |
| [42] | Webpage for animal monitoring using IoT and Cloud. | Behaviour | WSN, IoT, Cloud, GPS |
| [43] | Using Cloud and IoT for livestock monitoring. | Environment, health, growth, behavior, reproduction, emotions, and stress | GPS, Cloud, IoT |
| [44] | Mobile application using Cloud/edge computing; Meta data for digital certificates. | Location, disease | Cloud, IoT, digital certificates |
| [45] | Scalable IoT platforms integrated with cloud; tested against a batch of pigs at their fattening stage. | Feed monitoring | IoT, Cloud |
| [46] | Disease detection in poultry chickens using IoT. | Movement | IoT, Machine Learning |
| [47] | Monitoring livestock using IoT. | Temperature, humidity, light intensity | WSN, ZigBee, GPS |
| [48] | Using IoT and LoRa technologies for monitoring animal health. | Temperature, heart rate, humidity | IoT, LoRa, RFID |
| [49] | Poultry information management using cloud. | Temperature, humidity | WSN, Cloud |
| [50] | Review of poultry farming using IoT. | Vocal, heart rate | IoT, Artificial Intelligence, Infrared |
| [51] | Animal health diagnosis using Cloud. | Temperature, rumination time, and exercise | Cloud |
